# Mitofusins *Mfn1* and *Mfn2* are required to preserve glucose-but not incretin-stimulated beta cell connectivity and insulin secretion

**DOI:** 10.1101/2020.04.22.055384

**Authors:** Eleni Georgiadou, Charanya Muralidharan, Michelle Martinez, Pauline Chabosseau, Alejandra Tomas, Fiona Yong Su Wern, Elina Akalestou, Theodoros Stylianides, Asger Wretlind, Cristina Legido-Quigley, Ben Jones, Livia Lopez Noriega, Yanwen Xu, Guoqiang Gu, Nour Alsabeeh, Céline Cruciani-Guglielmacci, Christophe Magnan, Mark Ibberson, Isabelle Leclerc, Yusuf Ali, Scott A. Soleimanpour, Amelia K. Linnemann, Tristan A. Rodriguez, Guy A. Rutter

## Abstract

**Aims/hypothesis:** Mitochondrial glucose metabolism is essential for stimulated insulin release from pancreatic beta cells. Whether mitochondrial networks may be important for glucose or incretin sensing has yet to be determined.

**Methods:** Here, we generated mice with beta cell-selective, adult-restricted deletion of the mitofusin genes *Mfn1* and *Mfn2* (β*Mfn1/2* dKO). Whole or dissociated pancreatic islets were used for live beta cell fluorescence imaging of cytosolic or mitochondrial Ca^2+^ concentration and ATP production or GSIS in response to increasing glucose concentrations or GLP-1 receptor agonists. Serum and blood samples were collected to examine oral and i.p. glucose tolerance.

**Results:** β*Mfn1/2* dKO mice displayed elevated fed and fasted glycaemia (p<0.01, p<0.001) and a >five-fold decrease (p<0.0001) in plasma insulin. Mitochondrial length, glucose-induced polarisation, ATP synthesis and cytosolic Ca^2+^ increases were all reduced (p<0.05,p<0.01,p<0.0001) in dKO islets, and beta cell Ca^2+^ dynamics were suppressed *in vivo* (p<0.001). In contrast, oral glucose tolerance was near normal in β*Mfn1/2* dKO mice (p<0.05, p<0.01) and GLP-1 or GIP receptor agonists largely corrected defective GSIS from isolated islets through an EPAC-dependent signalling activation.

**Conclusions/interpretation:** Mitochondrial fusion and fission cycles are thus essential in the beta cell to maintain normal glucose, but not incretin, sensing. Defects in these cycles in some forms of diabetes might therefore provide opportunities for novel incretin-based or other therapies.

**Graphical abstract:** Impact of Mfn1/2 deletion on glucose and incretin stimulated-insulin secretion in beta cells. (A) In control animals, glucose is taken up by beta cells through GLUT2 and metabolised by mitochondria (elongated structure) through the citrate (TCA) cycle, leading to an increased mitochondrial proton motive force (hyperpolarised Δψm), accelerated ATP synthesis and O2 consumption rate (OCR). Consequently, the cytoplasmic ATP:ADP ratio rises, which causes closure of KATP channels, depolarisation of plasma membrane potential (ψm), opening of VDCCs and influx of cytosolic Ca^2+^. Elevated [Ca^2+^]cyt triggers a number of ATP-dependent processes including insulin secretion and improved beta-beta cell communication through connexin 36 (Cx36). (B) Following *Mfn1/2* deletion (β*Mfn1/2* dKO), highly fragmented mitochondria were associated with reduced mitochondrial Ca^2+^ ([Ca^2+^]m) accumulation, leading to a less polarised Δψm, weaker OCR, lower mtDNA copy number and decreased ATP synthesis. This is expected to result in weaker ψm depolarisation, cytosolic Ca^2+^ influx and beta-beta cell connectivity due to lower expression of Cx36. Despite observing a higher number of docked insulin granules on the plasma membrane, insulin secretion was highly suppressed in these animals. This was also associated with increased beta cell death and reduced beta cell mass. (C) In response to incretins, insulin secretion is potentiated through the activation of GLP1-R and cAMP signalling involving PKA- and EPAC2-dependent pathways. Elevated [Ca^2+^]cyt triggers a number of ATP-dependent processes including insulin secretion and Ca2+ removal into the endoplasmic reticulum (ER).(D) In β*Mfn1/2* dKO cells, activation of the GLP1-R was shown to be linked with a potentiation of the EPAC2 pathway that is PKA independent, along with an increased ER Ca^2+^ uptake and improved beta-beta cell communication. How these ‘amplifying’ signals of glucose metabolism for insulin secretion are linked with fragmented mitochondria remains unknown. Red and bold arrows represent enhanced pathways; dashed arrows represent impaired pathways. This figure was produced using illustrations from Servier Medical Art, http://smart.servier.com/

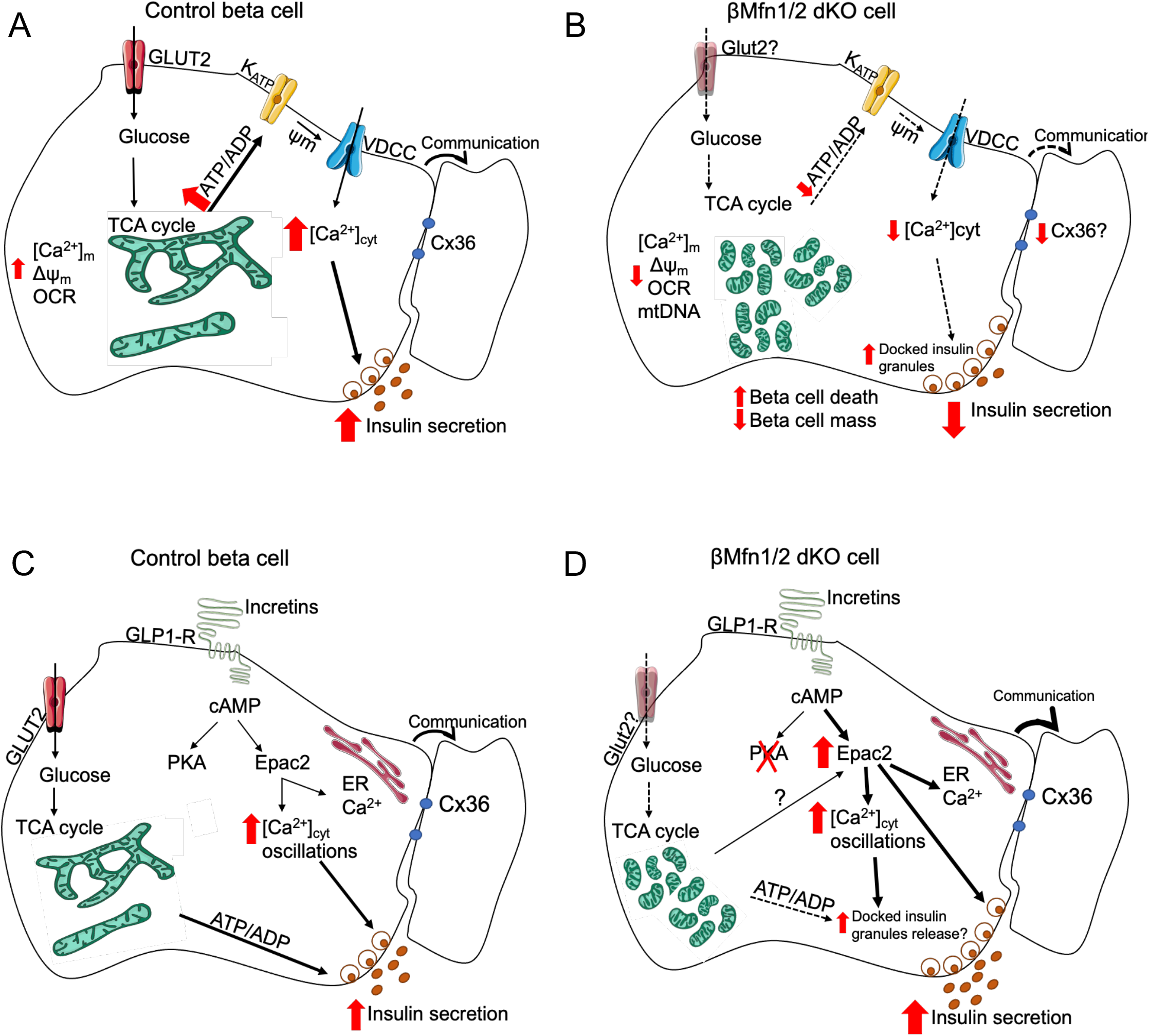

**Research in context:** **What is already known about this subject?**

Mitochondrial ultrastructural variations and number are altered in beta cells of human T2D patients [1].

Mice lacking *Opa1*, which controls mitochondrial fusion and inner membrane cristae structure, in beta cells, develop hyperglycaemia and defects in GSIS [2].

**What is the key question?**

Is an interconnected mitochondrial network essential in primary mouse beta cells for normal insulin secretion and glucose homeostasis?

**What are the new findings?**

We generated mice with beta cell-selective, adult-restricted deletion of the mitofusin genes *Mfn1* and *Mfn2* and show that insulin secretion and glucose homeostasis are strongly reduced *in vivo*.

Cytosolic and mitochondrial Ca^2+^ increases, Δψ_m_, ATP production and beta cell connectivity are impaired in β*Mfn1/2* dKO animals.

Incretins bypass the above defects through an exchange protein directly activated by cAMP (EPAC)-dependent signalling mechanism.

**How might this impact on clinical practice in the foreseeable future?**

The ability of incretins to bypass defects in mitochondrial function might be exploited by the design of new agonists which target this pathway.

## Introduction

Mitochondria are often referred to as the powerhouses or “chief executive organelles” of the cell, using fuels to provide most of the energy required to sustain normal function [3]. Mitochondrial oxidative metabolism plays a pivotal role in the response of pancreatic beta cells to stimulation by glucose and other nutrients [4]. Thus, as blood glucose increases, enhanced glycolytic flux and oxidative metabolism lead to an increase in ATP synthesis, initiating a cascade of events which involve the closure of ATP-sensitive K^+^ (K_ATP_) channels [5], plasma membrane depolarisation and the influx of Ca^2+^ via voltage-dependent Ca^2+^ channels (VDCC). The latter, along with other, less well defined “amplifying” signals [6], drive the biphasic release of insulin [4]. Gut-derived incretin hormones including glucagon-like peptide-1 (GLP-1) and glucose-dependent insulinotropic peptide (GIP) [7] further potentiate secretion by binding to class-B G-protein coupled receptors (GPCRs) to generate cAMP and other intracellular signals [8].

Changes in mitochondrial function in beta cells may also contribute to declining insulin secretion and to T2D [9] and has been described in several models of the disease [10]. Additionally, variants in mitochondrial DNA (mtDNA) in human populations are associated with altered T2D risk [11]. In animal models, alterations in beta cell mtDNA lead to reduced GSIS, hyperglycaemia and beta cell death [12].

Under normal physiological conditions, mitochondria undergo fusion and fission cycles which are essential for quality control and adaptation to energetic demands [13]. Thus, highly inter-connected mitochondrial networks allow communication and interchange of contents between mitochondrial compartments, as well as with other organelles such as the endoplasmic reticulum (ER) [14]. These networks exist interchangeably with more fragmented structures, displaying more “classical” mitochondrial morphology [15]. Mitochondrial fission is necessary for “quality control” and the elimination of damaged mitochondria by mitophagy [16].

Whilst the mitofusins MFN1 and MFN2, homologues of the *D. melanogaster* fuzzy onions (*fzo*) and mitofusin (*dmfn*) gene products [17], are GTPases that mediate fusion of the outer mitochondrial membrane (OMM), optic atrophy protein 1 (OPA1) controls that of the inner mitochondrial membrane (IMM). Dynamin related protein 1 (DRP1) is responsible for mitochondrial fission [18]. Other regulators include FIS1, mitochondrial fission factor (MFF) and MiD49/51 [19].

Changes in mitochondrial fusion and fission dynamics are observed in the pancreatic beta cell in animal models of diabetes [9, 20], and patients with T2D and obesity exhibit smaller and swollen mitochondria in pancreatic tissue samples [1]. Additionally, toxic islet amyloid polypeptide (IAPP) oligomers, usually co-expressed with insulin in beta cells, were present in both ER and mitochondrial membranes of T2D patients and rodents transgenic for human-IAPP (h-IAPP) [21].

Here, we explore the potential impact of mitochondrial fragmentation in the control of insulin secretion. We show that this manoeuvre exerts profound effects on insulin release, glucose homeostasis and Ca^2+^ dynamics. Remarkably, the deficiencies in insulin secretion are largely corrected by incretin hormones, suggesting a possible approach to ameliorating the consequences of mitochondrial fragmentation in some forms of diabetes.

## Methods

### Study approval

C57BL/6J mice were housed in individually ventilated cages in a pathogen-free facility at 22°C with a 10-14 h light-dark cycle and were fed *ad libitum* with a standard mouse chow diet (Research Diets, New Brunswick, NJ, USA). All *in vivo* procedures were approved by the UK Home Office, according to the Animals (Scientific Procedures) Act 1986 with local ethical committee approval under personal project license (PPL) number PA03F7F07 to I.L.

### Generation of beta cell selective *Mfn1/Mfn2* knockout (β*Mfn1/2* dKO), *Clec16a* null and Pdx1CreER mice

Animals were purchased and genotyped as described in ESM Methods.

### mRNA extraction and quantitative reverse transcription PCR

For measurements of mRNA levels, pancreatic islets from control and β*Mfn1/2* dKO mice were isolated by collagenase digestion [22]. Total RNA from islets (50-100) was extracted and reverse transcribed as previously described [23] (see ESM Table 2 for primer details).

### Tissue DNA extraction and measurement of mitochondrial DNA (mtDNA) copy number

Total islet DNA was isolated using Puregene Cell and Tissue Kit (Qiagen, Manchester, UK). See ESM Methods for further details.

### SDS-PAGE and western blotting

Islets were collected and lysed (20 μg) as previously described [23]. See ESM Methods for details.

### Intraperitoneal (i.p.) or oral gavage of glucose followed by insulin or ketone levels measurement and insulin tolerance assessment *in vivo*

IPGTTs, IPIITTs, OGTTs and plasma insulin measurements were performed as previously described [23] in control and β*Mfn1/2* dKO mice. β-ketones were measured in tail vein blood from fed or fasted (16h) mice using an Area 2K device (GlucoMen, Berkshire, UK).

### *In vitro* insulin secretion

Isolated islets were subjected to glucose-stimulated insulin secretion as described in ESM Methods.

### cAMP assay

Total cAMP was measured in primary dispersed mouse islet cells as described in ESM Methods.

### Single-cell fluorescence imaging

Pancreatic islets were isolated from mice, dissociated into single beta cells and plated onto glass coverslips [24]. See ESM Methods for details.

### Mitochondrial shape analysis

To determine morphological characteristics of mitochondria, confocal stacks were analysed with ImageJ using an in-house macro (available upon request). See ESM Methods for details.

### Whole-islet fluorescence imaging

Fluorescence imaging of whole islets was performed as described in ESM Methods.

### TIRF fluorescence imaging

Experiments using the membrane-located zinc sensor ZIMIR [25] or the genetically-encoded and vesicle-located green marker NPY-Venus were performed as presented in ESM Methods.

### Pancreas immunohistochemistry

Isolated pancreata were fixed and visualised as described in ESM Methods.

### Metabolomics/lipidomics

Plasma samples from control and dKO mice were analysed as described in ESM Methods.

### Measurement of oxygen consumption rate

Seahorse XF96 extracellular flux analyzer (Seahorse Bioscience, Agilent, Santa Clara, CA, USA) was used for intact mouse islets respirometry as described in ESM Methods.

### Electron microscopy (EM)

**Fixed** islets were processed as described in ESM Methods.

### *In vivo* Ca^2+^ imaging of AAV8-INS-GCaMP6s infected endogenous pancreatic islets

Pancreatic islets of control and β*Mfn1/2*-KO mice were imaged *in vivo* as described in ESM Methods.

### Connectivity analysis

#### Pearson (*r*)-based connectivity and correlation analyses

Correlation analyses were performed as described in ESM Methods.

### Monte Carlo-based signal binarisation and data shuffling for identification of highly connected cells

Data were analysed using approaches as previously described [26, 27]. For further details see ESM Methods.

### RNA-Seq data analysis

Processing and differential expression analysis of RNA-Seq data was performed as in [28] and ESM Methods.

### Statistics

Data are expressed as mean ± SEM unless otherwise stated. Significance was tested by Student’s two-tailed t-test and Mann–Whitney correction or two-way ANOVA with Sidak’s multiple comparison test for comparison of more than two groups, using GraphPad Prism 8 software (San Diego, CA, USA). p< 0.05 was considered significant. Experiments were not randomised or blinded.

## Results

### Generation of a conditional β*Mfn1/2* dKO mouse line

Efficient deletion of *Mfn1* and *Mfn2* in the beta cell was achieved in adult mice using the Pdx1-Cre^ERT2^ transgene and tamoxifen injection at 7-8 weeks. Possession of this transgene alone had no effect on glycaemic phenotype or cellular composition of pancreatic islets (Suppl. Fig.1**A-C**). Deletion of mitofusin genes was confirmed by qRT-PCR (Fig.1A) and Western (immuno-) blotting (Fig.1B) analysis in 14-weeks old male mice. Relative to *β-actin*, expression of the *Mfn1* and *Mfn2* transcripts in isolated islets from dKO mice decreased by ∼83 and 86% accordingly vs control islets (Fig.1A; p<0.01,p<0.0001), consistent with selective deletion in the beta cell compartment [29]. No differences were detected in the expression of other mitochondrial fission and fusion mediator genes such as *Opa1*, *Drp1* and *Fis1* (Fig.1A). Body weights also differed between groups after 20-21 weeks (Suppl.Fig.**2A;** p<0.05).

**Fig. 1.**
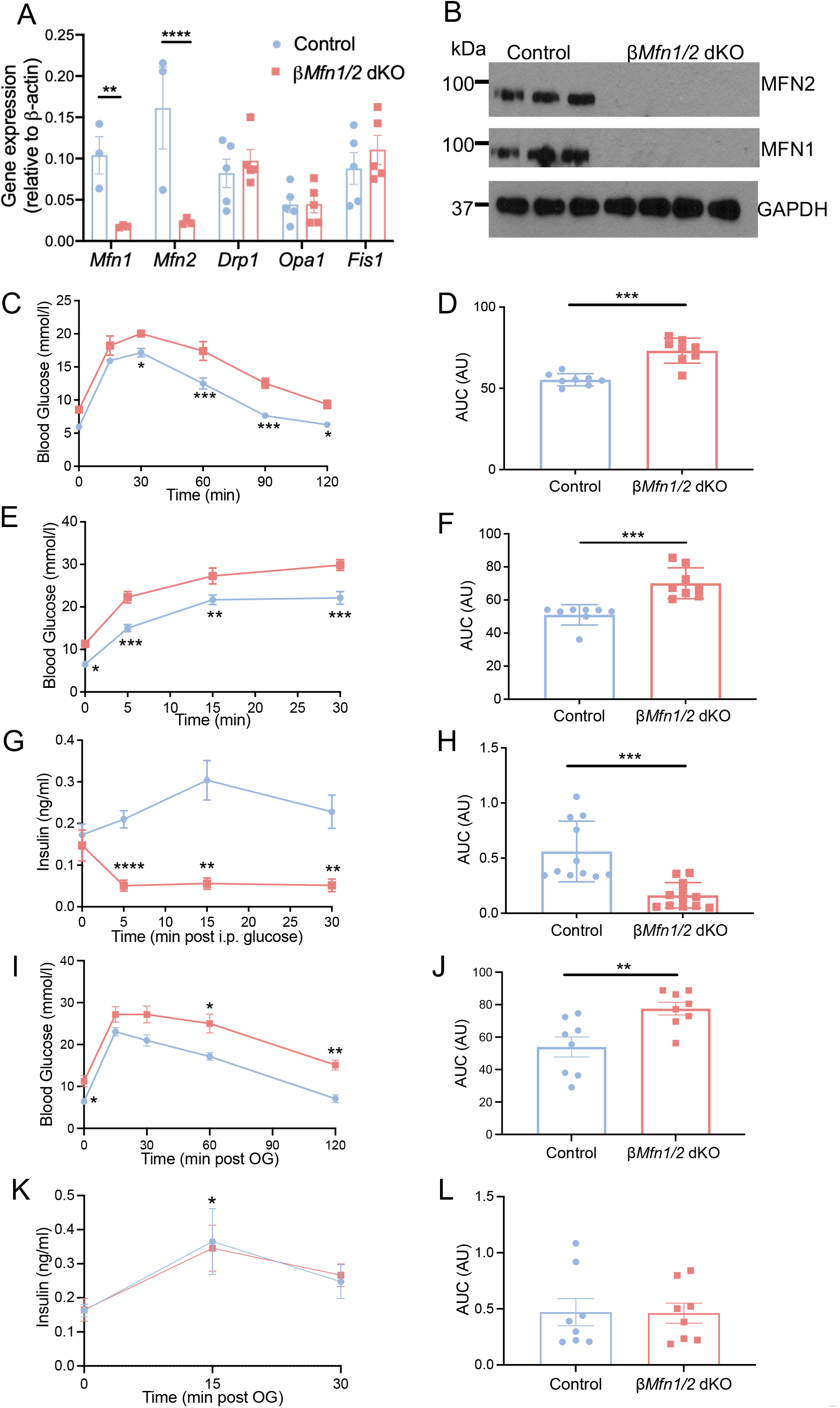
Generation of a conditional β*Mfn1/2* dKO mouse line which displays a highly impaired glucose tolerance *in vivo*. (A) qRT-PCR quantification of *Mfn1, Mfn2, Drp1, Opa1* and *Fis1* expression in control and dKO islets relative to *β-actin* (*n=*3-5 mice per genotype in two independent experiments).(B) Western blot analysis demonstrating efficient MFN1 (84 kDa) and MFN2 (86 kDa) deletion relative to GAPDH (36 kDa) in isolated islets (*n*=3-4 mice per genotype in three independent experiments).(C) Glucose tolerance was measured in dKO mice and littermate controls by IPGTT (1 g/kg body weight).(D) Corresponding AUC from (C) (*n*=8 mice per genotype,in 2 independent experiments). (E) Glucose tolerance measured by IPGTT (using 3 g/kg body weight) and (F) the corresponding AUC were assessed in β*Mfn1/2* dKO and control mice (*n*=8 mice per genotype in two independent experiments). (G) Plasma insulin levels during IPGTT in dKO and control mice (*n*=11-12 mice per genotype in three independent experiments) and (H) the corresponding AUC. (I) Glucose tolerance post-oral gavage (3 g/kg body weight) was measured in *n*=8 animals per genotype in two independent experiements.The corresponding AUC is shown in (J). (K) Plasma insulin levels during OGTT in dKO and control mice (*n*=8 mice per genotype in two independent experiments) and (L) the corresponding AUC. (Blue, *control* mice; red, *dKO* mice. Data are presented as mean±SEM. *p<0.05; **p<0.01; ***p<0.001; ****p<0.0001 as indicated, or *control* vs *dKO* mice at the time points as indicated in (**K**), analysed by unpaired two-tailed Student’s t-test and Mann– Whitney correction or two-way ANOVA test and Sidak’s multiple comparisons test. All experiments were performed in 14-week-old male mice.

### β*Mfn1/2* dKO mice are glucose intolerant with impaired GSIS *in vivo*

To study the effects of mitofusin gene deletion in beta cells on systemic glucose homeostasis and insulin secretion *in vivo,* i.p. injections (IPGTT) were performed on β*Mfn1/2* dKO and control mice (Fig.1C). Glucose challenge revealed impaired glucose tolerance in dKO mice compared to their control littermates with levels of glucose being higher at most time points following glucose injection (Fig.1C**-D**;p<0.05,p<0.001). Glucose intolerance was even more prominent in 20-weeks old dKO mice (Suppl. Fig.2**B-C**; p<0.001, p<0.0001). β*Mfn1/2* dKO mice (with a 27 mmol/l glycaemia at 15 min.; Fig.1E**-F**;p<0.05; p<0.01; p<0.001) showed dramatically lower insulin levels upon glucose challenge vs control animals, indicating a severe insulin secretory deficiency (Fig.1G-H,p<0.01,p<0.001,p<0.0001). In contrast, following an oral gavage (Fig.1I-J), the plasma insulin levels in dKO mice (with a 27 mmol/l glycaemia at 15min.) were indistinguishable from control animals (Fig.1K-L;p<0.05; 0 vs 15min. in dKO). Insulin tolerance was unaltered insulin tolerance in β*Mfn1/2* dKO mice vs control littermates (Suppl.Fig.**1D-E**). Nevertheless, dKO mice displayed significantly elevated plasma glucose (Suppl.Fig.**1F)** under both fed and fasted conditions. Additionally, an increase in β-ketones (ketone bodies) was observed in fasted dKO vs control mice (Suppl.Fig.**1G**). These changes were inversely related to plasma insulin levels, which were lower in dKO than control mice under both fed and fasted, conditions (Suppl.Fig.**1H**).

### Deletion of *Mfn1/2* alters mitochondrial morphology in beta cells

Mitochondrial morphology was assessed using confocal imaging and digital deconvolution. Mitochondria were elongated in dissociated control beta cells (Fig.2A) while the mitochondrial network in dKO cells was highly fragmented (Fig.2A; and inset). The number of mitochondria per cell was not altered (Fig.2B). Mitochondrial elongation and perimeter were significantly decreased in β*Mfn1/2* dKO cells, while circularity on the other hand, was increased indicative of rounder and smaller organelles (Fig.2B; p<0.0001). Mitochondrial structure was also evaluated in isolated islets by transmission electron microscopy (TEM), confirming the presence of more highly fragmented mitochondria in dKO mouse islets compared to the control group (Fig.2C). Cristae structure and organisation were also markedly altered in β*Mfn1/2* dKO islet cells (Fig.2C;enlarged panels and schematic representations).

**Fig. 2.**
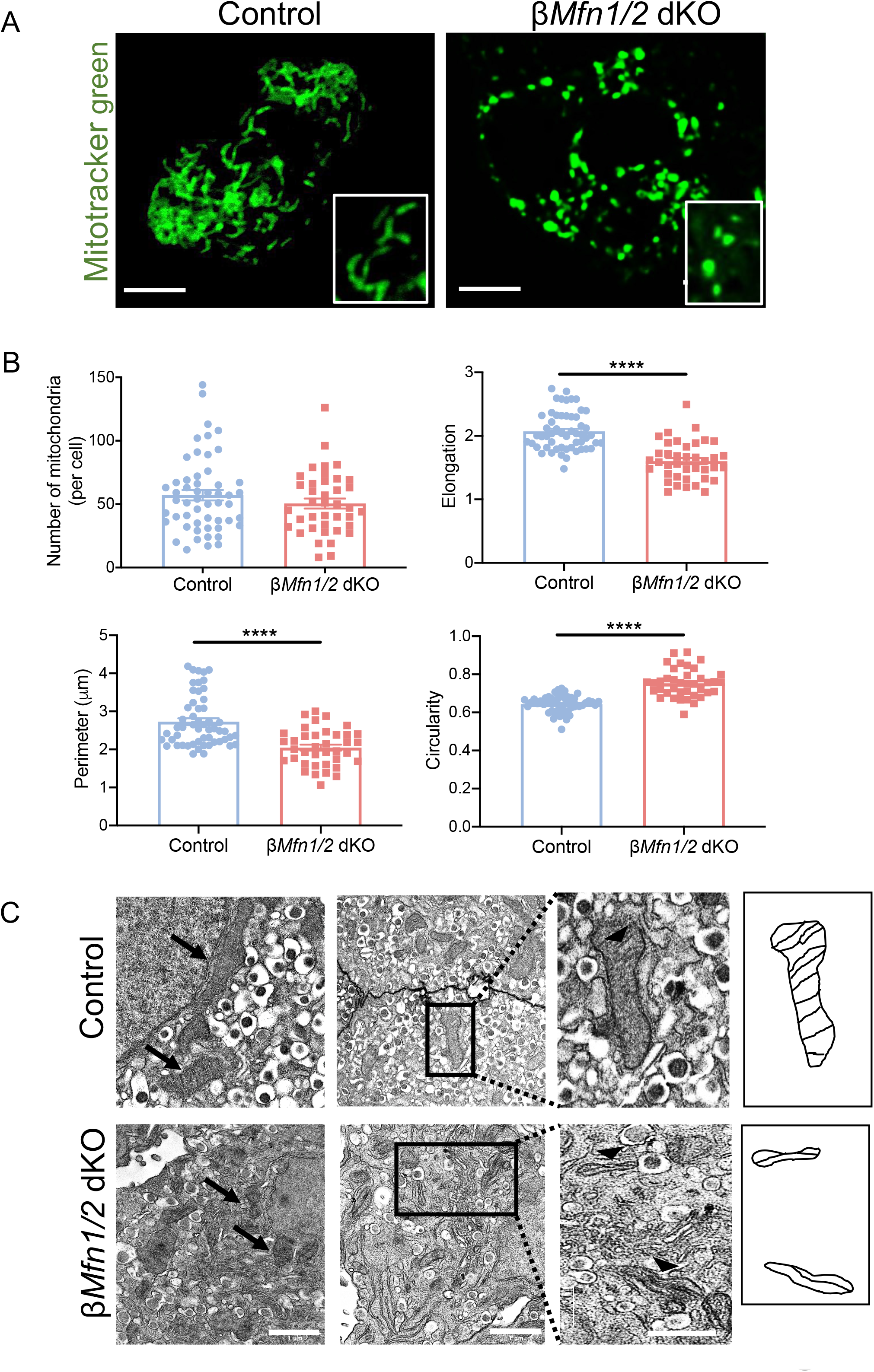
Mitochondrial ultrastructure is altered following *Mfn1/2* deletion. (A) Confocal images of the mitochondrial network of dissociated beta cells stained with Mitotracker green; scale bar: 5 μm. Lower right panels: magnification of selected areas. (B) Mitochondrial morphology analysis on deconvolved confocal images of dissociated beta cells. A macro was developed to quantify the number of mitochondria per cell and measure the elongation, perimeter and circularity (0: elongated; 1: circular mitochondria) of the organelles in control and dKO animals (*n*=40-54 cells; *n*=3 mice per genotype). (C) Electron micrographs of mitochondria indicated with black arrows in islets isolated from control and dKO mice; scale bars: 1μm. Right panel: magnification of selected areas showing the cristae structure (black arrow heads); scale bar: 0.5 μm. Schematic representation of enlarged mitochondria. Data are presented as mean±SEM. ****p<0.0001 as indicated, analysed by unpaired two-tailed Student’s t-test and Mann–Whitney correction. Experiments were performed in 14-week-old male mice.

### *Mitofusin* deletion leads to modest changes in beta cell mass

Immunohistochemical analysis of pancreata from dKO mice showed a small but significant (∼33%) loss of pancreatic beta (insulin-positive) cells vs the control group (Fig.3A-B; p<0.05). Alpha (glucagon-positive) cell surface was not affected by the loss of mitofusin genes (Fig.3C). However, *Mfn1* and *Mfn2* loss was associated with a ∼53% reduction in beta cell-alpha cell ratio (Fig.3D; p<0.05). In line with these findings, the number of TUNEL-positive beta cells were markedly increased in dKO vs control animals (Fig.3E-F;p<0.05), suggesting that programmed cell death contributes to the observed decrease in beta cell mass.

**Fig. 3.**
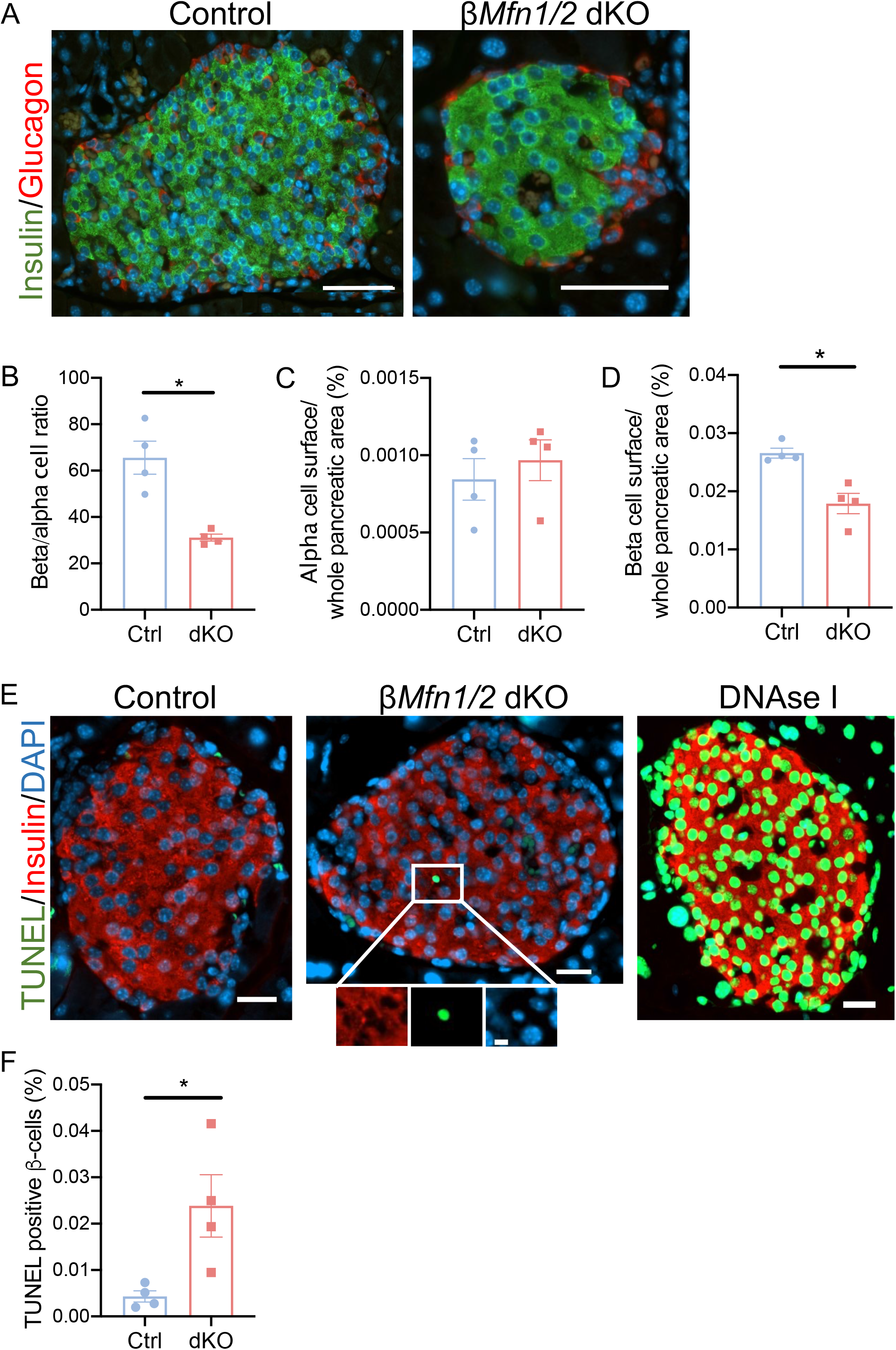
Absence of *Mfn1/2* in beta cells leads to decreased beta cell mass and increased beta cell apoptosis. (A) Representative pancreatic sections immunostained with glucagon (red) and insulin (green); scale bars: 50μm.(B) The beta cell and alpha cell surface (C) measured within the whole pancreatic area in control and dKO mice were determined, as well as the beta/alpha cell ratio in (D), (*n*=79-86 islets, 4 mice per genotype; experiment performed in triplicate).(E) Representative confocal images of islets with TUNEL positive (green) apoptotic beta cells (ROI) and insulin (red). Magnification of selected area displaying each fluorescent channel; scale bar: 5μm. DNase I treated sections were used as a positive control in the TUNEL assay. Scale bars: 20μm.(F) Quantification of the percentage of islets containing TUNEL positive cells (*n*=114-133 islets, 4 mice per genotype; experiment performed in triplicate). Data are presented as mean±SEM. *p<0.05, assessed by unpaired two-tailed Student’s t-test and Mann–Whitney correction. Experiments were performed in 14-week-old male mice.

### Beta cell identity is modestly altered in β*Mfn1/2* dKO islets

Whilst *Ins2*, *Ucn3* and *Glut2* (*Slc2a2*) were significantly downregulated, *Trpm5* was upregulated, in dKO islets (Suppl.Fig.**3**). No changes in α- or beta cell disallowed genes [30] were detected. In contrast, genes involved in mitochondrial function such as *Smdt1* and *Vdac3* were upregulated in β*Mfn1/2* dKO islets, consistent with compromised mitochondrial Ca^2+^ uptake, and ATP production, respectively, in dKO beta cells (Suppl.Fig.**3**). Lastly, genes involved in ER stress and mito/autophagy were also affected by inactivation of *Mfn1* and *Mfn2* with *Chop (Ddit3)* and *p62* being upregulated and *Lc3* and *Cathepsin L* downregulated.

### Glucose-induced cytosolic Ca^2+^ and Δψ_m_ changes are impaired in β*Mfn1/2* dKO beta cells *in vivo*

Mitochondrial membrane polarisation (Δψ_m_) and Ca^2+^ dynamics were next studied *in vivo*. Animals previously infected with GCaMP6s, and co-stained with tetramethyl rhodamine methyl ester (TMRM) immediately prior to data capture, were imaged for 18-30 min. post i.p. injection of glucose in control (Fig.4A; Suppl.Fig.**4A**) and dKO mice (Fig.4B; Suppl.Fig.**4B**). Imaging revealed cytosolic Ca^2+^ oscillations ([Ca^2+^]_cyt_; upward traces) and synchronous mitochondrial membrane depolarisation (downward traces; TMRM positive organelles) in response to elevated glucose in control beta cells. These were largely abolished in dKO islets in response to glucose (Fig.4B;Suppl.Fig.**4B**). Measurement of the AUC of fold change traces above baseline depicted significantly impaired GCaMP6s spike signals in response to glucose (Fig.4C;p<0.001 and p<0.05; Suppl.Fig.**4C**) and a tendency towards less TMRM uptake in dKO islets (Fig.4D).

**Fig. 4.**
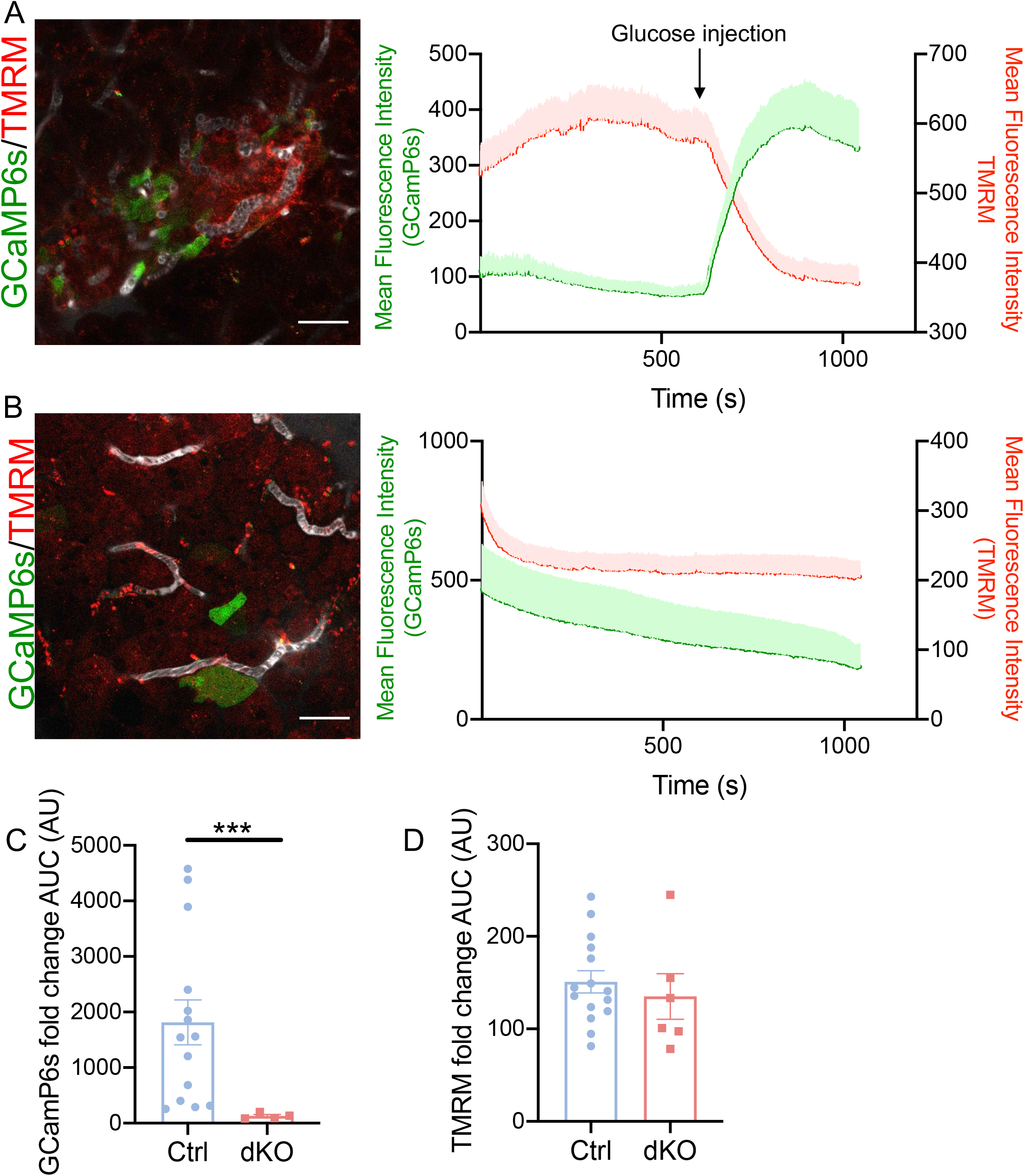
Deletion of Mfn1/2 impairs beta cell function *in vivo*. Representative *in vivo* images of GCaMP6s labelled islets and TMRM stained mitochondria surrounded by their vasculature in control and dKO mice. (A) Representative traces depicting fluorescence intensity of cytosolic Ca^2+^ (GCaMP6s) and mitochondrial TMRM signals in control (ESM Video 1) and (B) dKO animals (ESM Video 2) before and after glucose injection as indicated; scale bars: 45 µm; (*n*=2 animals per genotype). (C) AUC of fold change measurements above baseline for each GCaMP6s and (D) TMRM traces measured (*n*=4-15 total responding cells). Under these conditions, glucose concentrations in control mice were 17.1±2.5 mmol/l and 32.1±3.9 mmol/l in dKO animals after glucose injection. Analysis was performed on the most responsive beta cells where oscillations could be detected in both groups.Green, GCaMP6s; red, TMRM signals. Data are presented as mean±SEM. ***p<0.001, assessed by unpaired two-tailed Student’s t-test and Mann–Whitney correction. Experiments were performed in 20-week-old male mice.

### Mitofusins are essential to maintain normal glucose-stimulated Ca^2+^ dynamics, mitochondrial membrane potential and ATP synthesis in beta cells

Increased cytosolic Ca^2+^ is a major trigger of insulin exocytosis in response to high glucose [4]. dKO mouse islets exhibited a significantly lower increase in [Ca^2+^]_cyt_ compared to control islets (Fig.5A-C;p<0.01). When the K_ATP_ channel opener diazoxide and a depolarising K^+^ concentration (20 mmol/l KCl) were then deployed together to bypass the regulation of these channels by glucose, cytosolic Ca^2+^ increases were not significantly impaired in dKO compared to control animals (Fig.5B-C). A substantial reduction in mitochondrial free Ca^2+^ concentration ([Ca^2+^]_mito_) in response to 17 mmol/l glucose [23, 31] was also observed in dKO islets (Fig.5D-F; p<0.05). Of note, subsequent hyperpolarisation of the plasma membrane with diazoxide caused the expected lowering of mitochondrial [Ca^2+^]_mito_ in control islets (reflecting the decrease in [Ca^2+^]_cyt_;**Fig.5E-F**), but was almost without effect on dKO islets.

**Fig. 5.**
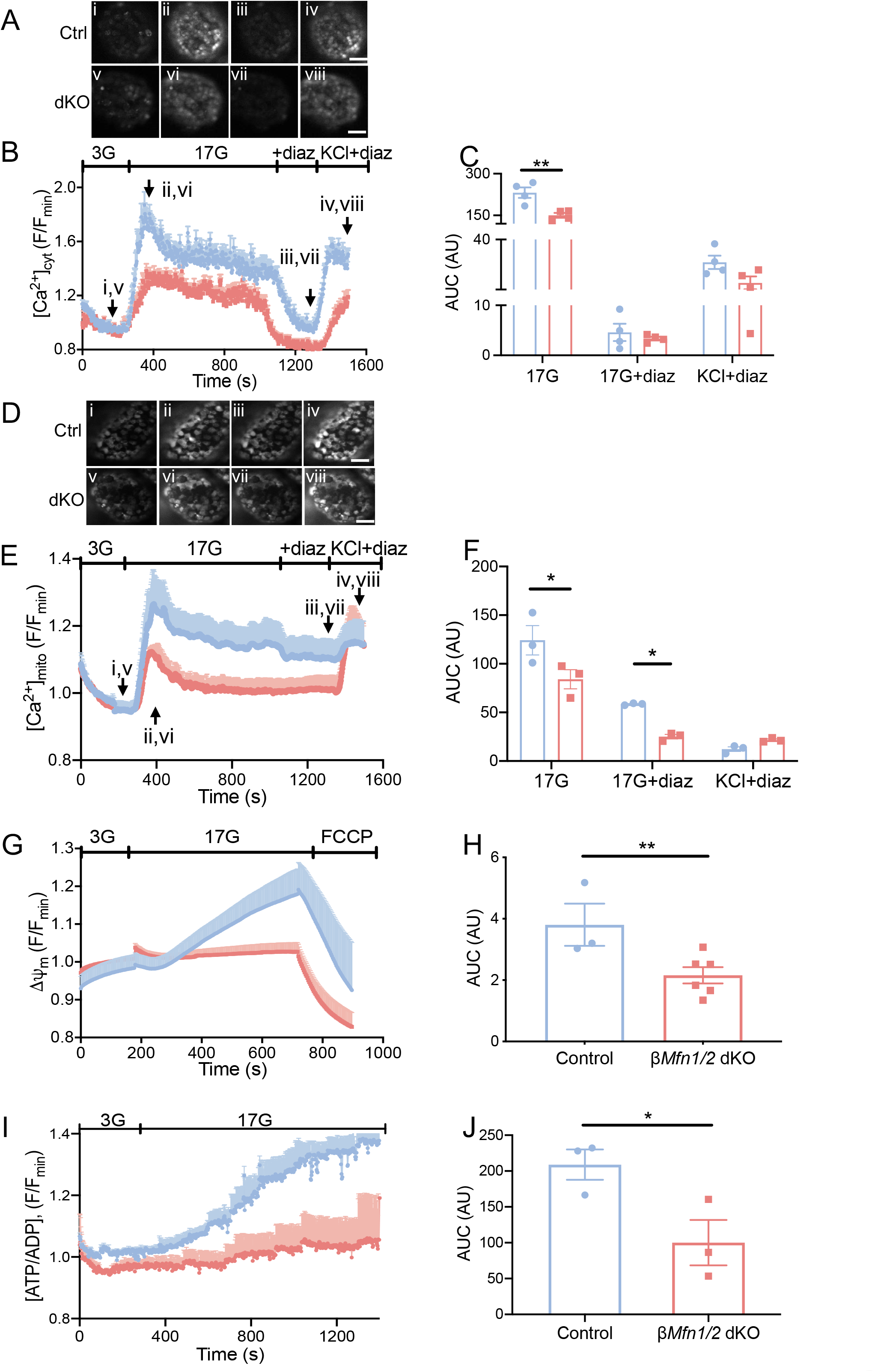
*Mfn1/2* deletion from pancreatic beta cells impairs cytosolic and mitochondrial Ca^2+^ uptake and changes mitochondrial potential and ATP synthesis *in vitro*. (A) Each snapshot of isolated control (i–iv) and dKO-derived (v–viii) islets was taken during the time points indicated by the respective arrows in (B). Scale bar: 50 μm. See also ESM Video 3. (B) [Ca^2+^]_cyt_ traces in response to 3G, 3 mmol/l glucose, 17 mmol/l glucose (17G; with or without diazoxide [diaz]) or 20 mmol/l KCl with diaz were assessed following Cal-520 uptake in whole islets. Traces represent mean normalised fluorescence intensity over time (F/F_min_).(C) The corresponding AUC is also presented (*n=*17-26 islets, 4 mice per genotype); 17G AUC measured between 245 s and 1045 s, 17G+diaz AUC measured between 1200 s and 1320 s, and KCl+diaz AUC measured between 1424 s and 1500 s. For each genotype different baselines (ctrl diaz/KCl: 0.95, dKO diaz/KCl: 0.8 were taken into consideration to measure AUCs.(D) Each snapshot of isolated control (i–iv) and dKO-derived (v–viii) islets was taken during the time points indicated by the respective arrows in (E). Scale bar: 50 μm. See also ESM Video 4. (E) [Ca^2+^]_mito_ changes in response to 17G (with or without diazoxide [diaz]) and 20 mmol/l KCl were assessed in islets following R-GECO infection. Traces represent mean normalised fluorescence intensity over time (F/F_min_).(F) The corresponding AUC is also shown (*n*=20-23 islets, 3 mice *per* genotype; 17G AUC measured between 270 s and 1100 s, 17G+diaz AUC measured between 1101 s and 1365 s and KCl AUC measured between 1366 s and1500 s).(G) Dissociated beta cells were loaded with TMRE to measure changes in Δψ_m_, and perifused with 3 mmol/l glucose (3G), 17G or FCCP as indicated. Traces represent normalised fluorescence intensity over time (F/Fmin).(H) AUC was measured between 700–730 s (under 17G exposure) from the data shown in (G) (*n*=146-254 cells,3-6 mice per genotype).(I) Changes in the cytoplasmic ATP:ADP ratio ([ATP:ADP]) in response to 17 mmol/l glucose (17G) was examined in whole islets using the ATP sensor Perceval.(J) AUC values corresponding to (I) were measured between 418– 1400 s (under 17G exposure) (data points from *n*=22-23 islets, 3-6 mice per genotype). Data are presented as mean±SEM. *p<0.05, **p<0.01, assessed by unpaired two-tailed Student’s t-test and Mann–Whitney correction or two-way ANOVA test and Sidak’s multiple comparisons test. Experiments were performed in 14-week-old male mice.

Glucose-induced increases in Δψ_m_ were also sharply reduced in dKO vs control mouse islets (Fig.5G-H; p<0.01). Addition of 2-[2-[4-(trifluoromethoxy)phenyl]hydrazinylidene]-propanedinitrile (FCCP) resulted in a similar collapse in apparent Δψ_m_ in islets from both genotypes (Fig.5G). To assess whether mitochondrial fragmentation may impact glucose-induced increases in mitochondrial ATP synthesis we performed real-time fluorescence imaging using Perceval (Fig.5I-J). While control islets responded with a time-dependent rise in the ATP:ADP ratio in response to a step increase in glucose from 3 mmol/l to 17 mmol/l, β*Mfn1/2* dKO beta cells failed to mount any response (Fig.5J;p< 0.05).

### Beta cell-beta cell connectivity is impaired by *Mfn1/2* ablation

Intercellular connectivity is required in the islet for a full insulin secretory response to glucose [10, 26]. To assess this, individual Ca^2+^ traces recorded from Cal-520-loaded beta-cells in mouse islets (Fig.5A-B) were subjected to correlation (Pearson *r*) analysis to map cell-cell connectivity (Suppl.Fig.**5A**). Following perfusion at 17 mmol/l glucose, β*Mfn1/2* dKO beta cells tended to display an inferior, though not significantly different, coordinated activity than control cells, as assessed by counting the number of coordinated cell pairs (Suppl.Fig.**5C**; 0.94 vs 0.90 for control vs dKO, respectively). By contrast, beta cells displayed highly coordinated Ca^2+^ responses upon addition of 20 mmol/l KCl in dKO islets. Similarly, analysis of correlation strength in the same islets revealed significant differences in response to 17 mmol/l glucose between genotypes. In fact, dKO islets had weaker mean beta-beta cell coordinated activity (Suppl**. Fig.5B, D**; p<0.05; 0.88 vs 0.77 for control vs dKO, respectively), indicating that mitofusins affect the strength of connection rather than the number of coordinated beta cell pairs. A tending towards lower expression of the gap junction gene *Cx36/Gjd2* has also been observed in dKO islets (Suppl.Fig.**5E**).

Clear adherence to a power law distribution of connected beta cells [26, 27] was apparent in the control islet group in the elevated glucose condition where 5.70% of the beta cells hosted at least 60% of the connections with the rest of the beta cells (‘hubs’;Suppl.Fig.6; R^2^=0.15). No clear adherence to a power-law distribution of connected beta cells was present in the dKO group (R^2^=0.002) despite displaying a higher percentage (15.06%) of beta cell-beta cell connections.

### Unaltered ER Ca^2+^ mobilisation but decreased mitochondrial O_2_ consumption and mtDNA depletion in β*Mfn1/2* dKO islets

No differences in cytosolic Ca^2+^ responses between genotypes were observed after agonism at the Gq-coupled metabotropic acetylcholine (Ach) receptor (Fig.6A-C). In contrast, measurements of O_2_ consumption revealed that both basal and glucose-stimulated mitochondrial respiratory capacities were significantly impaired in dKO islets (Fig.6D-E). Moreover, dKO islets displayed a ∼75% reduction in mtDNA (Fig.6F;p<0.05).

**Fig. 6.**
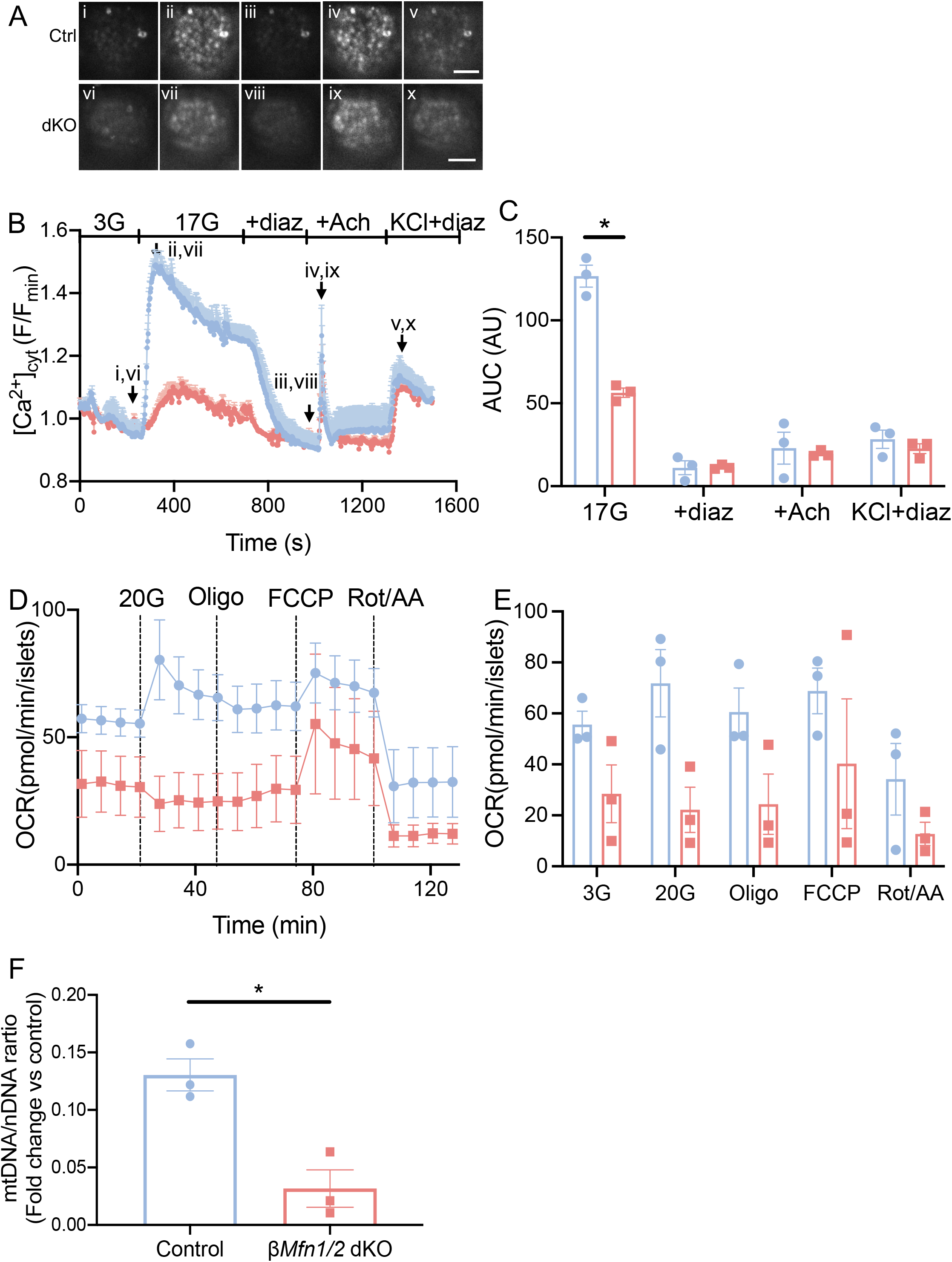
O_2_ consumption and mtDNA are deleteriously affected when *Mfn1/2* are abolished in beta cells, while [Ca^2+^]_ER_ mobilisation remains unchanged. (A) Each snapshot of isolated control (i–v) and dKO-derived (vi–x) islets was taken during the time points indicated by the respective arrows in (B). Scale bar: 50 μm. See also ESM Video 5. (B) Changes in [Ca^2+^]_ER_ were measured in whole islets incubated with Cal-520 and perifused with 17 mmol/l glucose (17G; with or without diazoxide [diaz]), 17G with 0.1 mmol/l acetylcholine (Ach) and diaz, or 20 mmol/l KCl with diaz (C) AUC values corresponding to (B) were measured (17G AUC measured between 260 s and 740 s, 17G+diaz AUC measured between 846 s and 1020 s, 17G+diaz+Ach AUC measured between 1021 s and 1300 s and KCl AUC measured between 1301 s and 1500 s) (n=29-31 islets, 3 mice per genotype). (D) Representative oxygen consumption rate (OCR) traces of islets (∼10 per well) were acutely exposed to 20 mmol/l glucose (final concentration), Oligomycin A (Oligo), FCCP, and Rotenone with Antimycin A (AA) (performed in triplicate, in two independent experiments).(E) Average values for each condition corresponding to (D).(F) The relative mitochondrial DNA copy number was measured by determining the ratio of the mtDNA-encoded gene *mt-Nd1* to the nuclear gene *Ndufv1* (*n*=3 mice per genotype).Data are presented as mean±SEM. *p<0.05, assessed by unpaired two-tailed Student’s t-test and Mann– Whitney correction or two-way ANOVA test and Sidak’s multiple comparisons test. Experiments were performed in 14-week-old male mice.

### Impaired GSIS *in vitro* and beta cell connectivity can be rescued by incretins in β*Mfn1/2* dKO mouse islets

While GSIS was markedly impaired in dKO islets (Fig.7A;p<0.05), incretins (GLP-1 or GIP), or the GLP1R agonist exendin-4, at a submaximal concentration of 10 mmol/l glucose, led to a significant potentiation in GSIS in both groups (control: 3G vs ex4; p<0.05 and dKO: 3G vs ex4; p<0.0001, or 3G vs GLP-1; p<0.001, or 3G vs GIP; p<0.001). Consequently, insulin secretion in response to 10 mmol/l glucose was no longer different between control and β*Mfn1/2* dKO islets after incretin addition (Fig.7A-B). Moreover, under these conditions, forced increases in intracellular cAMP imposed by the addition of FSK or IBMX, which activate adenylate cyclase (AC) and inhibit phosphodiesterase (PDE) respectively, also eliminated differences in GSIS between the genotypes (Fig.7B). No differences in insulin secretion were observed between control and dKO islets after depolarisation with KCl.

**Fig. 7.**
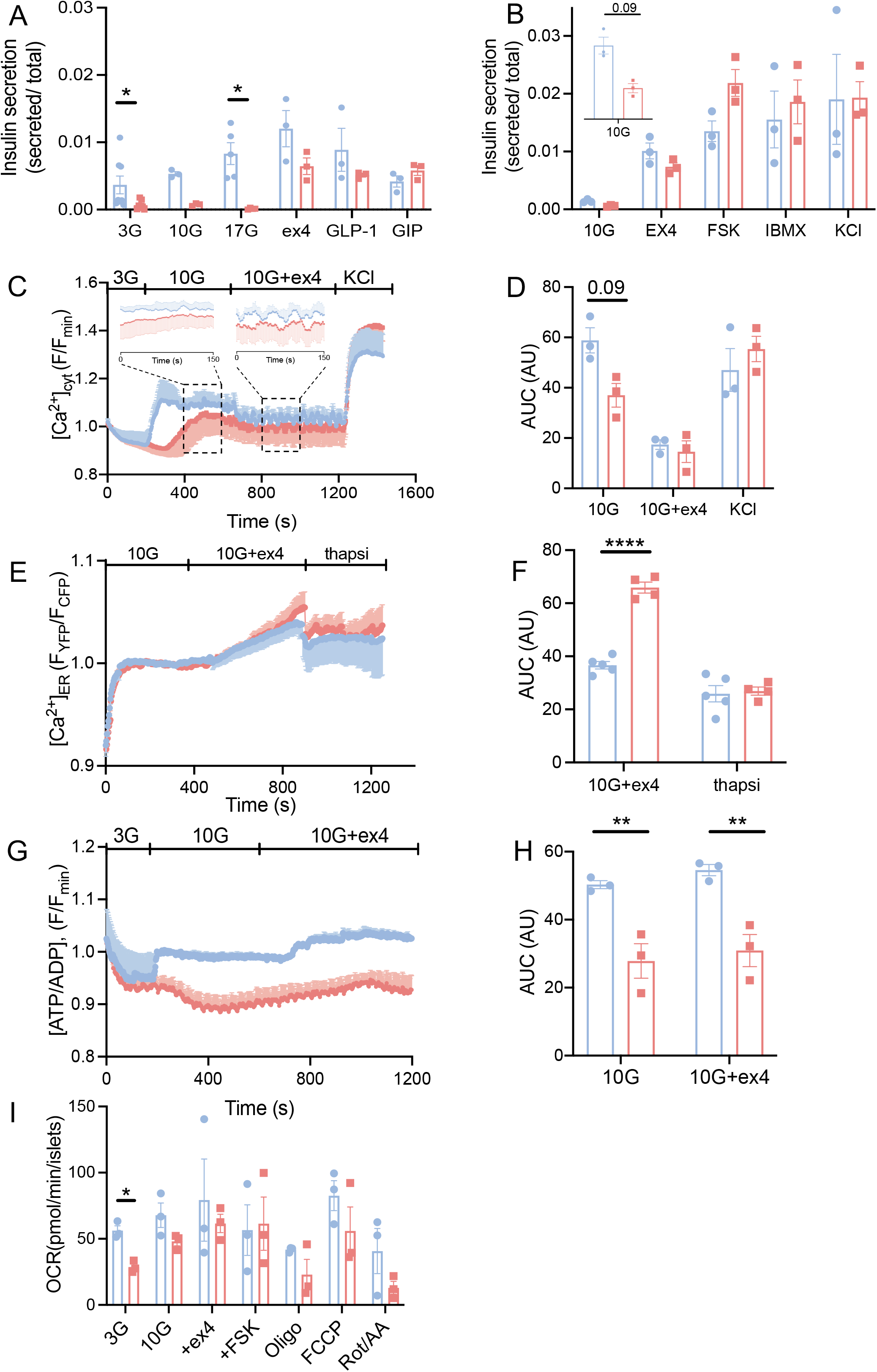
Impaired insulin secretion can be rescued by GLP-1R agonists *in vitro* by increasing cytosolic Ca^2+^ oscillation frequency. (A) Insulin secretion measured during serial incubations in batches in 3 mmol/l glucose (3G), 10 mmol/l glucose (10G), or 100 nmol/l exendin-4 (ex4), GLP-1 or GIP in presence of 10G or 17 mmol/l glucose (17G) (*n=*4-7 mice per genotype in two independent experiments).(B) Insulin secretion measured during serial incubations in batches in 10 mmol/l glucose (10G), supplemented with 100 nmol/l exendin-4 (ex4), 10 µmol/l FSK or 100 µmol/l IBMX and 20 mmol/l KCl (*n=*3 mice per genotype, in two independent experiments). (C) [Ca^2+^]_cyt_ changes in response to 3G, 3 mmol/l glucose, 10 mmol/l glucose (10G; with or without exendin-4 [ex4]) or 20 mmol/l KCl were assessed following Cal-520 uptake in whole islets. Traces represent mean normalised fluorescence intensity over time (F/F_min_). See also ESM video 6. Dashed ROIs represent fluorescent segments of extended time scales. Both control and dKO traces reveal faster oscillatory frequencies in response to exendin-4. (D) The corresponding AUC is also presented (*n=*19-20 islets, 3 mice per genotype; 10G AUC measured between 200 s and 660 s, 10G+ex4 AUC measured between 800 s and 950 s), and KCl AUC measured between 1200 s and 1500 s).(E) Dissociated beta cells were loaded with D4ER to measure changes in [Ca^2+^]_ER_, and perifused with 10 mmol/l glucose (10G), 10G+ex4 or thapsigargin (10G+thapsi) as indicated. Traces represent corrected ratio values post-linear fitting over time. (F) AUC was measured between 350–900 s (under 10G+ex4) and 900-1300 s (10G+thapsi) from the data shown in (E) (*n*=44-46 cells,4-5 mice per genotype). (G) Changes in cytoplasmic ATP:ADP ratio ([ATP:ADP]) in response to 10G or 10G with 100nmol/l ex4 was examined in whole islets.(H) AUC values corresponding to (G) were measured between 185-720s (under 10G exposure) or 721-1200s (under 10G with ex4) (data points from *n*=3 mice per genotype). (I) Average OCR values of islets (∼10 per well) that were exposed to 3mmol/l or 10mmol/l glucose (final concentration), 10mmol/l glucose supplemented with ex4, FSK, Oligomycin A (Oligo), FCCP, and Rotenone with Antimycin A (AA) (*n*=3 mice per genotype; experiment performed in duplicate). Data are presented as mean±SEM. *p<0.05;**p<0.01, ****p<0.0001 assessed by two-way ANOVA test and Sidak’s multiple comparisons test. Experiments were performed in 14-week-old male mice.

We next explored whether the incretin-mediated improvements in insulin secretion in response to incretins were the result of altered [Ca^2+^]_cyt_ dynamics. Islets from isolated dKO mice displayed a delayed increase in [Ca^2+^]_cyt_ in response to 10 mmol/l glucose compared to control islets (Fig.7C-D; p=0.09; AUC control: 10G vs ex4; p<0.05; dKO: 10G vs ex4; p<0.001; ex4 vs KCl; p<0.05). Addition of exendin-4 led to the emergence of oscillatory activity in both groups and under these conditions, differences between genotypes, as seen in Fig.5B, were no longer evident (Fig.7C). Measured at 10mmol/l glucose, control and dKO islets displayed increases in ER Ca^2+^ in response to exendin-4 (Fig.7E-F; AUC;p<0.001) while the response exaggerated in the latter group. Neither group displayed significant changes in ATP:ADP ratio in response to exendin-4 (Fig.7G-H;AUC;p<0.01). Analysis of OCR revealed no significant differences between genotypes at 10mmol/l glucose in the presence or absence of exendin-4 or FSK (Fig.7I; p<0.05). Finally, exendin-4 sharply increased beta cell-beta cell connectivity in dKO, but not in control islets, as assessed by monitoring Ca^2+^ dynamics and the number of correlated cell pairs (Fig. 8A,C) or Pearson *r* value (Fig. 8B,D).

**Fig. 8.**
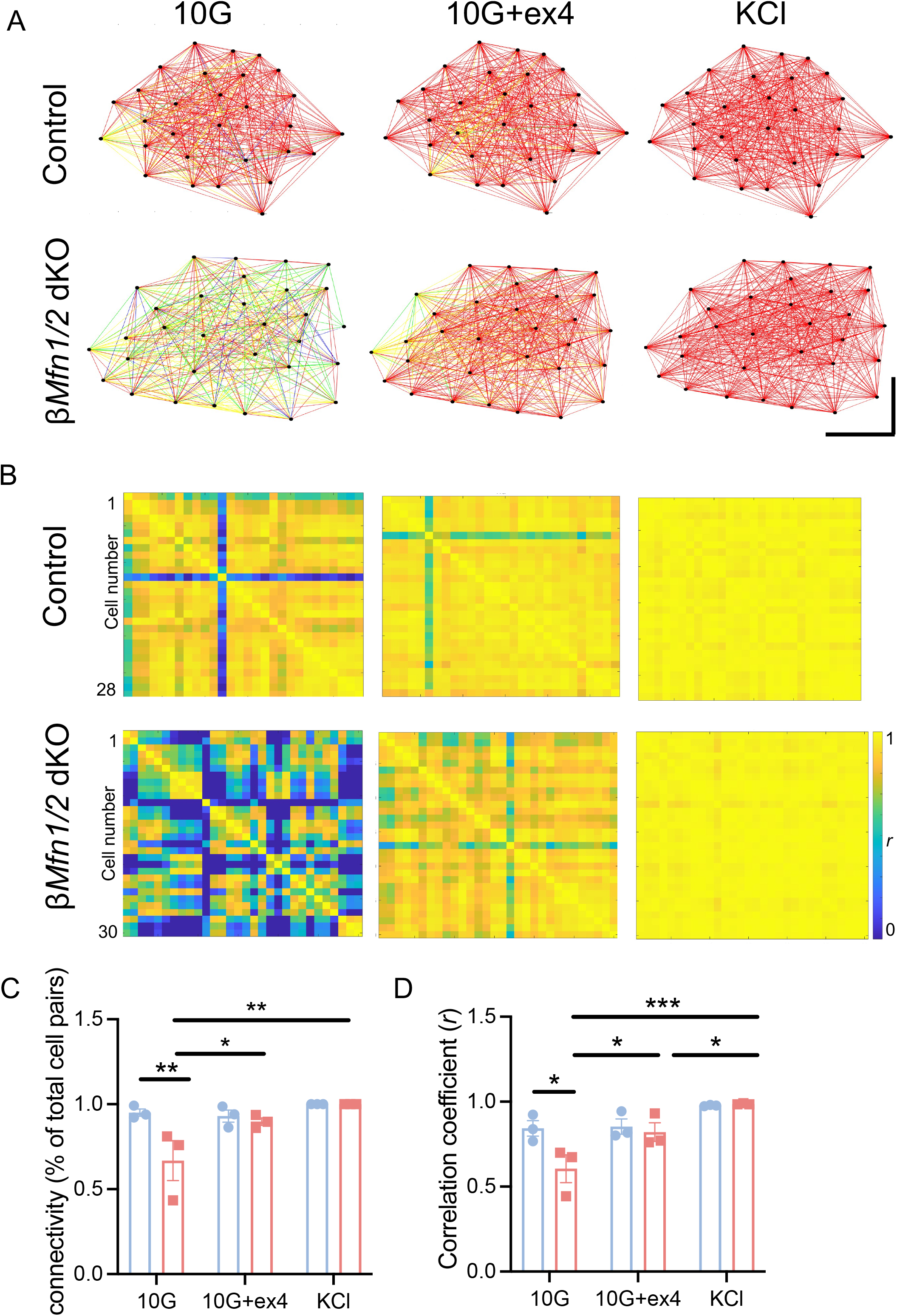
The GLP1-R agonist, exendin-4, improves intercellular connectivity in βMfn1/2 dKO β-cells. (A) Representative cartesian maps of control and dKO islets with colour coded lines connecting cells according to the strength of Pearson analysis (colour coded *r* values from 0 to 1, blue to red respectively) under 10mmol/L (10G), 10mmol/L with 100nmol/l exendin-4 (10G+ex4) glucose or 20mmol/L KCl; scale bars: 40 μm.(B) Representative heatmaps depicting connectivity strength (*r*) of all cell pairs according to the colour coded *r* values from 0 to 1, blue to yellow respectively.(C) Percentage of connected cell pairs at 10G, 10G+ex4 or KCl (*n*=19-20 islets, 3 mice per genotype).(D) *r* values between β-cells in response to glucose, exendin-4 or KCl (*n*=3 mice per genotype).Data are presented as mean±SEM. *p<0.05,**p<0.01, ***p<0.001 assessed by two-way ANOVA test and Sidak’s multiple comparisons test. Experiments were performed in 14-week-old male mice.

### Insulin secretion is rescued by incretins through an EPAC-dependent activation

Neither basal nor incretin-stimulated cAMP levels differed between control and dKO groups (Fig.9A). Nonetheless, insulin secretion was amplified in the presence of the protein kinase A (PKA) inhibitor H89 alone or in addition to IBMX and FSK in dKO islets (Fig.9B; p<0.05, p<0.01, p<0.001). Insulin secretion was further increased in dKO islets when EPAC was selectively activated, while PKA was inhibited by H89 (Fig.9C; p<0.05).

**Fig. 9.**
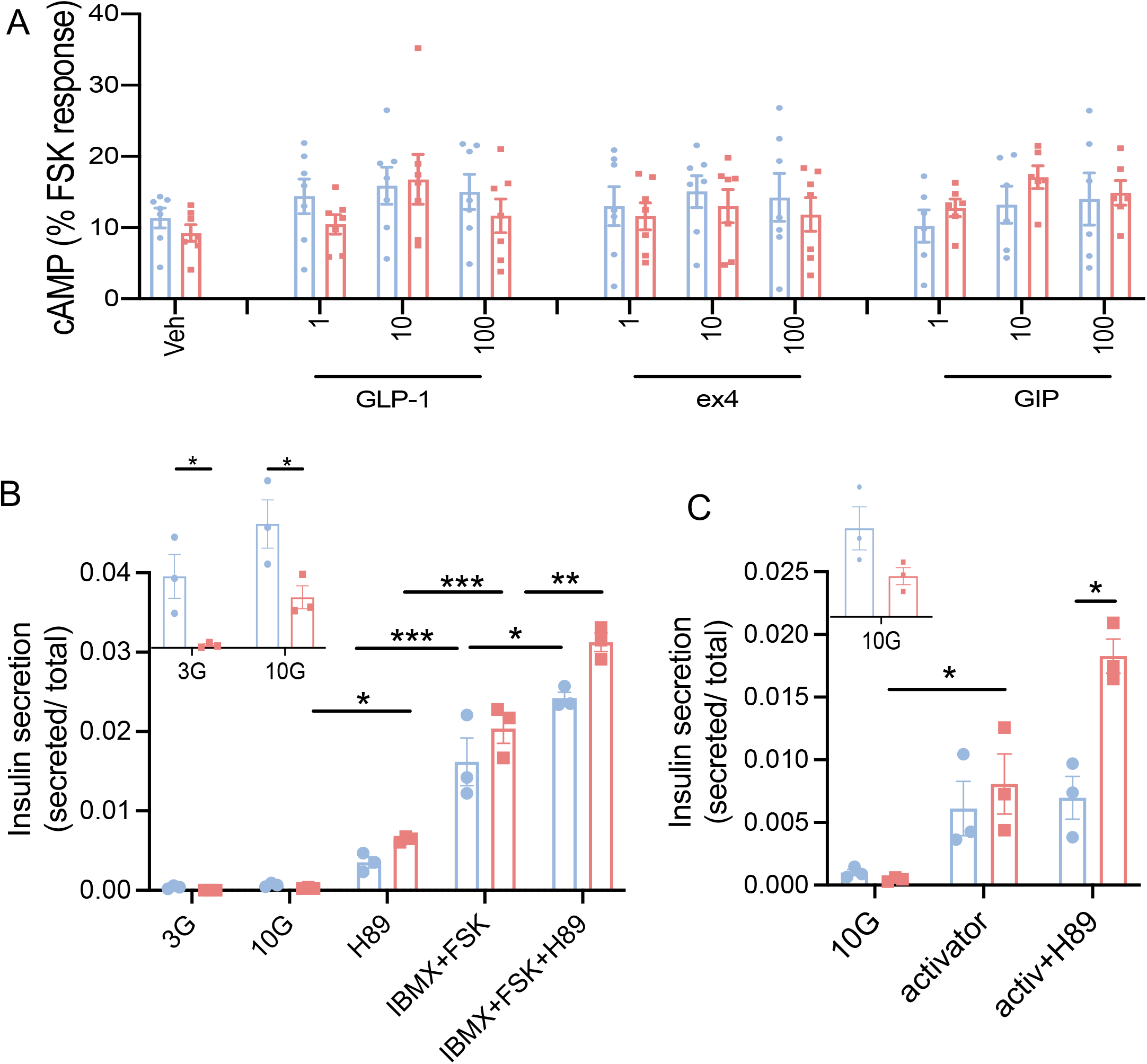
Insulin secretion is rescued through an EPAC-dependent activation in dKO islets. (A) Concentration of cAMP (normalised to FSK) in response to 1,10,100 nmol/l GLP-1, ex4 and GIP stimulation in dissociated islets (*n*=6-7 animals, in 2 independent experiments). (B) Insulin secretion measured during serial incubations in batches in 3 mmol/l glucose (3G), 10 mmol/l glucose (10G), or 10 mmol/l glucose supplemented with 10µmol/l H89, 10 µmol/l FSK with 100 µmol/l IBMX or H89 (*n=*3 mice per genotype, in two independent experiments). (C) Insulin secretion measured during serial incubations in batches in 10 mmol/l glucose (10G), or 10 mmol/l glucose supplemented with 6µmol/l EPAC-activator, or EPAC-activator with 10µmol/l H89 (*n=*3 mice per genotype, in two independent experiments). Data are presented as mean±SEM. *p<0.05,**p<0.01, ***p<0.001 assessed by two-way ANOVA test and Sidak’s multiple comparisons test. Experiments were performed in 14-week-old male mice.

### Defective glucose-stimulated insulin secretion is rescued by GLP-1R agonism in *Clec16a* null mice

To determine whether incretins may reverse defective insulin secretion in an alternative model of mitochondrial dysfunction, we examined mice lacking the mitophagy regulator *Clec16a* selectively in the pancreatic islet (*Clec16a*^Δpanc^) [32]. Glucose-stimulated insulin secretion was sharply inhibited in null vs Pdx1-Cre control mice, and these differences between genotype were largely corrected in by the addition of exendin-4 (Suppl. Fig.**7A**; p<0.0001). Correspondingly, whereas the difference between *Clec16a*^Δpanc^ and control mice was significant for IPGTTs there was no such (significant) difference for the OGTTs at 15mins, in line with the findings above for β*Mfn1/2* dKO mice (Suppl. Fig.**7B-C**; p<0.05, p<0.01).

### Defective secretion of a preserved pool of morphologically-docked granules in β*Mfn1/2* dKO mouse beta cells

To determine whether the markedly weaker stimulation of insulin secretion in dKO islets may reflect failed recruitment of secretory granules into a readily releasable or morphologically-docked pool beneath the plasma membrane, we next deployed total internal reflection fluorescence (TIRF) microscopy in dissociated beta cells. By over-expressing the secretory vesicle marker neuropeptide Y-Venus (NPY-Venus), the number of insulin granules was significantly higher in close proximity with the plasma membrane in dKO cells after treatment with 20 mmol/l KCl (Suppl.Fig.**8A-B;**p<0.05). However, when we then used the cell surface-targeted Zinc indicator to monitor induced exocytotic release (ZIMIR) [25] in response to depolarisation as a surrogate for insulin secretion, release events were fewer in number and smaller in dKO (Suppl.Fig.**8C-E**).

### Altered plasma metabolomic and lipidomic profiles in β*Mfn1/2* dKO mice

We applied an -omics approach to study metabolite and lipid changes in peripheral plasma samples from control and dKO mice (Suppl.Fig.**9**). Of 29 metabolites, the levels of five metabolic species (shown in red) were significantly altered in β*Mfn1/2* dKO compared to control animals (Suppl.Fig.**9A;** p<0.05;p<0.01). In the lipidomics analysis, 298 lipid species from 17 different classes were studied. When comparing dKO to control samples, the majority of lipid classes displayed a remarkably homogeneous downward trend of the individual lipid species they comprised (Suppl.Fig.**9B;** p<0.05;p<0.01).

### Changes in *Mfn1* and *Mfn2* expression in mouse strains maintained on regular chow or high fat high sugar (HFHS) diet

To determine whether the expression of *Mfn1* or *Mfn2* might be affected under conditions of hyperglycaemia mimicking T2D in humans, we interrogated data from a previous report [33] in which RNA sequencing was performed on six mouse strains. BALB/cJ mice showed “antiparallel” changes in *Mfn1* and *Mfn2* expression in response to maintenance on high fat high sugar (HFHS) diet for 10 days, and similar changes were obtained in DBA/2J mice at 30 and 90 days (Suppl.Fig.**10A-B**).

## Discussion

The key goal of the present study was to determine the impact of disrupting mitochondrial dynamics on glucose- and incretin-stimulated insulin secretion. Deletion of both mitofusin isoforms selectively from the adult beta cell led to fragmentation of the mitochondrial network, impaired glucose signalling and altered beta cell identity. These changes were associated with marked dysglycaemia, which worsened with age. We chose the above strategy over the deletion of either mitofusin gene alone given the similar levels of expression of both in the beta cell [34] and the likelihood of at least partial functional redundancy as reported in [35]. Specifically, this recent report [35], using a complementary strategy (deletion with the constitutive Ins1*Cre* deleter strain), supports this view, demonstrating minor phenotypic effects of deletion in the beta cell of *Mfn1* or *Mfn2* alone.

The present study shows for the first time that collapse of the mitochondrial network prompted by the loss of *Mfn1* and *Mfn2* has a drastic impact on beta cell function. Our findings are in line with earlier studies which provided evidence for a critical role for preserved mitochondrial dynamics in insulin secretion [36, 37]. In these earlier studies, deletion of *Drp1* from primary mouse beta cells resulted in glucose intolerance, impaired GSIS and abnormal mitochondrial morphology. Conversely, over-expression of DRP1 in clonal INS1 cells decreased GSIS and increased the levels of apoptosis [38], suggesting that a balance between fission and fusion is critical to avoid pathological changes. Finally, mice deficient for *Opa1* in the beta cell develop hyperglycaemia, and show defects in the electron transport chain complex IV, Ca^2+^ dynamics, and insulin secretion [2]. None of the above studies explored the effects on incretin-stimulated secretion.

A striking finding in the present report is that, in contrast to results during intraperitoneal glucose injection, insulin secretion and glucose excursion were largely normal in dKO mice during OGTTs, where an incretin effect is preserved [7]. Incretins act, at least in large part, by increasing intracellular cAMP, and adequate cAMP levels are required for the normal stimulation of insulin secretion as glucose concentrations rise [7]. Conversely, incretin action requires adequate glucose levels (and hence intracellular metabolism of the sugar) [7]. Others [39, 40] have previously suggested that cAMP-raising agents may rescue the metabolic signalling defects associated with T2D. However, we are not aware of any previous studies which have *selectively* interfered with mitochondrial function, and then explored the ability of incretins to induce a reversal of the secretory deficiency.

What mechanisms might explain the ability of incretins to bypass defective insulin secretion after disruption of mitochondrial networks? Explored in dissociated islets, cAMP levels were not different between control and dKO animals in the absence or presence of hormones or GLP1R agonists. cAMP acts at multiple points in the secretory pathway, regulating plasma membrane excitability and Ca^2+^ dynamics (partly via exchange protein activated by cAMP and mainly EPAC2 translocation to granule docking sites on the plasma membrane) [40–42] and also on the exocytotic machinery via N-ethylmaleimide-sensitive attachment receptor (SNARE) proteins [43]. Here we show that insulin secretion stimulated by incretins is potentiated in dKO cells through an EPAC-dependent activation, while PKA has an inhibitory effect on secretion. It is still unknown how fragmented mitochondria are associated with the positive regulation of insulin secretion by EPAC and more studies need to be undertaken to explore this phenomenon. Synaptotagmin-7, another critical regulator of Ca^2+^-mediated exocytosis in beta cells, is known to be phosphorylated and activated by PKA [40], and may thus represent a potential target for incretin action in dKO cells. No difference in mobilization of Ca^2+^ was observed between groups using the IP3R agonist Ach, implying that ER Ca^2+^ stores are not depleted. Nevertheless, by mitofusin deletion, perifusion of exendin-4 revealed that Ca^2+^ accumulation in the ER was higher in dKO cells. This could be associated with a rapid mobilisation and increase in [Ca^2+^]_cyt_ oscillatory response that lead to an enriched pulsatile insulin secretion and beta cell connectivity [44]. Moreover, since [Ca^2+^]_m_ uptake was impaired in dKO cells and maintaining a high [Ca^2+^]_ER_ is important for beta cell function and survival, the ER could work as a rescue machinery to avoid [Ca^2+^]_cyt_ overload and toxicity or inflammatory stress [44].

Of note, impaired glucose-induced ATP synthesis and O_2_ consumption seen in fragmented dKO mitochondria were not recovered by incretins (at least under 10mmol/l glucose). This suggests that the acute rescue of insulin secretion is, to a substantial extent, independent of (and thus downstream to) changes in mitochondrial oxidative metabolism, in line with reports [45] that O_2_ consumption is only weakly affected by incretins in wild-type islets.

Importantly, we demonstrate that preserved mitochondrial ultra-structure is critical for normal beta cell-beta cell connectivity [27]. The mechanisms underlying these changes are, however, unclear but could be associated with altered *Cx36/Gjd2* expression, phosphorylation or activity, and hence the formation of gap junctions between beta cells [46]. Of note, highly connected “hub” [26] and leader [27] beta cell populations have been proposed to be particularly reliant on mitochondrial function [27]. Nevertheless, we did not observe any loss of hierarchical behaviour or apparent hub cell number in dKO mice.

Metabolomic analyses revealed that the dKO mouse provides a useful model of defective beta cell function observed, to differing extents, in both type 1 and type 2 diabetes [47–49].

Might changes in *Mfn1* and *Mfn2* expression be involved in diabetes development in rodents or humans? Changes in the expression of both genes were observed in two mouse models of the disease, in line with previous findings in beta cells overexpressing h-IAPP [21]. However, we are not aware of studies reporting changes in the expression of human *MFN1* or *MFN2* in human beta cells in this setting.

Our findings show that acute treatment with incretins, commonly used as treatments for T2D and obesity [7], largely reverses the deficiencies in insulin secretion and cytosolic Ca^2+^ signalling. We also demonstrate that highly selective impairments of mitochondrial function in beta cells can be rescued or bypassed by incretin treatment, and suggest that this might be an important mechanism of action for this drug class.

## Supporting information

All supplemental data

## List of abbreviations

[Ca^2+^]_cyt_: Cytoplasmic Ca^2+^ concentration
[Ca^2+^]_mito_: Mitochondrial free Ca^2+^ concentration
AA: Antimycin A
Ach: Acetylcholine
*Clec16a*^Δpanc^: Pancreatic islet specific Clec16a knock-out
Diaz: Diazoxide
dKO: double knock-out
Ex4: Exendin-4
FCCP: Carbonyl cyanide-4-phenylhydrazone
GIP: Glucose-dependent insulinotropic peptide
GLP-1: Glucagon-like peptide-1
GSIS: Glucose-stimulated insulin secretion
IMM: Inner mitochondria membrane
IPGTT: Intraperitoneal glucose tolerance test
OGTT: Oral gavage and glucose tolerance test
Oligo: Oligomycin
OMM: Outer mitochondrial membrane
r: Pearson correlation coefficient
Rot: Rotenone
TMRE: Tetramethylrhodamine ethyl ester
β*Mfn1/2* dKO: beta cell specific Mitofusin 1 and 2 double knock-out
Δψ_m_: Mitochondrial membrane potential

## Acknowledgements

We thank Stephen M. Rothery from the Facility for Imaging by Light Microscopy (FILM) at Imperial College London for support with confocal and widefield microscopy image recording and analysis. We also thank Aida Di Gregorio from the National Heart and Lung Institute (Imperial College) for genotyping the mice.

## Data Availability

Not applicable.

## Funding

GAR was supported by a Wellcome Trust Senior Investigator Award (098424AIA) and Investigator Award (212625/Z/18/Z), MRC Programme grants (MR/R022259/1, MR/J0003042/1, MR/L020149/1), an Experimental Challenge Grant (DIVA, MR/L02036X/1), an MRC grant (MR/N00275X/1), and a Diabetes UK grant (BDA/11/0004210, BDA/15/0005275, BDA16/0005485). IL was supported by a Diabetes UK project grant (16/0005485). This project has received funding from the Innovative Medicines Initiative 2 Joint Undertaking, under grant agreement no. 115881 (RHAPSODY). This Joint Undertaking receives support from the European Union’s Horizon 2020 research and innovation programme and EFPIA. This work is supported by the Swiss State Secretariat for Education, Research and Innovation (SERI), under contract no. 16.0097. AT was supported by MRC project grant MR/R010676/1. Intravital imaging was performed using resources and/or funding provided by National Institutes of Health grants R03 DK115990 (to AKL), Human Islet Research Network UC4 DK104162 (to AKL; RRID:SCR_014393). BJ acknowledges support from the Academy of Medical Sciences, Society for Endocrinology, The British Society for Neuroendocrinology, the European Federation for the Study of Diabetes, an EPSRC capital award and the MRC (MR/R010676/1). SAS was supported by the JDRF (CDA-2016-189, SRA-2018-539, COE-2019-861), the NIH (R01 DK108921, U01 DK127747), and the US Department of Veterans Affairs (I01 BX004444).

## Author contributions

EG performed experiments and analysed data. EG supported the completion of confocal and widefield microscopy and analysis. ATC performed the EM sample processing and data analysis. CM, MM and AKL were responsible for the *in vivo* intravital Ca^2+^ imaging in mice. PC contributed to the analysis and manipulation of the *in vivo* intravital Ca^2+^ measurements as well as the preparation and imaging of TIRF samples. TS contributed to the generation of the MATLAB script used for connectivity analysis. FYSW and YA generated and performed Monte Carlo-based signal binarisation. BJ performed the cAMP assays. EA and LLN performed the oral gavage in live animals. YX and GG performed studies with Pdx1CreER mice. NA assisted with Seahorse experiment protocols. CLQ and AW contributed to the metabolomics analysis. CCG, CM and MI were responsible for the RNAseq data analysis. SAS performed studies with Clec16a mice. TAR was involved in the design of the floxed *Mfn* alleles. TAR and IL were responsible for the maintenance of mouse colonies and final approval of the version to be published. GAR designed the study and wrote the manuscript with EG with input and final approval of the version to be published from all authors. GAR is the guarantor of this work and, as such, had full access to all the data in the study and takes responsibility for the integrity of the data and the accuracy of the data analysis.

## Declaration of interests

Authors’ relationships and activities GAR has received grant funding and consultancy fees from Les Laboratoires Servier and Sun Pharmaceuticals. The remaining authors declare that there are no relationships or activities that might bias, or be perceived to bias, their work.

## Electronic Supplemental Information

Supplemental Figures (1-10)

Supplemental Figure legends

## Electronic Supplemental Tables

**ESM Table 1.** Sequence of primers used for genotyping *Mfn1* and *Mfn2* flox.

**ESM Table 2.** List of primers used for qRT-PCR.

**ESM Table 3.** Metabolite differences found in plasma samples of control vs dKO mice according to metabolic class and both fold-change and t-test criteria.

**ESM videos 1-6**

## Notes

### Competing Interest Statement

GAR is a consultant for Sun Pharmaceuticals and has received grant support from Les Laboratories Servier

### Summary of Updates

Now included are new data on the signalling pathways through which incretins prompt insulin secretion after the disruption of Mfn1 and Mfn2, notably roles for Epac2. New controls are also included showing that the Pdx1CreERT deleter strain per se has no impact on glycemia, insulin secretion or the cellular composition of islets. N-values for IPGTTs have been increased

