## Supplementary material for "Mitofusins *Mfn1* and *Mfn2* are required to preserve glucose-but not incretin-stimulated beta cell connectivity and insulin secretion": All supplemental data

### ESM Methods

**Generation of beta cell-selective *Mfn1/Mfn2* knockout (beta *Mfn1/2* dKO), *Clec16a* null and Pdx1CreER mice** C57BL/6J male mice bearing *Mfn1* (Mfn1$^{tm2Dcc}$; JAX stock #026401) and *Mfn2* (B6.129(Cg)-Mfn2$^{tm3Dcc}$/J; JAX stock #026525; The Jackson Laboratory, Bar Harbor, ME,USA) alleles [1] with *lox*P sites flanking exons 4 and 6 were purchased from the Jackson laboratory and crossed to C57BL/6J transgenic animals carrying an inducible *Cre* recombinase under *Pdx1* promoter control (Pdx1-Cre$^{ERT2}$) [2]. Mice bearing floxed *Mfn* alleles but lacking *Cre* recombinase were used as littermate controls in this study. Mice were genotyped following protocols described by the Jackson laboratory for each of these strains. (See ESM Table 1 for genotyping primer details). Recombination was achieved by daily tamoxifen (20mg/ml [diluted in corn oil; Sigma-Aldrich, Dorset, UK]) i.p. injections for five days at 7-8 weeks of age in both control and β*Mfn1/2* dKO (dKO) groups. Body weight was monitored every week, for 14 weeks, in *ad libitum* fed mice. Unless otherwise stated, all experiments were performed on 14-weeks aged male mice.

Animals with floxed *Clec16a* alleles were bred to mice carrying the Pdx1-Cre transgene, resulting in efficient deletion in the islet after tamoxifen treatment (*Clec16a*$^{\Delta panc}$) as previously described [3].Pdx1-Cre alone mice were used as littermate controls.

Pdx1CreER mice were generated as previously described [2]. Tamoxifen [4mg; Sigma-Aldrich, Dorset, UK] was injected i.p three times at alternative days in 4-weeks old mice. IPGTTs (2 g/kg) glucose) and pancreatic tissues were examined 4 weeks after tamoxifen injection.

**Tissue DNA extraction and measurement of mitochondrial DNA (mtDNA) copy number** Total islet DNA was isolated using Puregene Cell and Tissue Kit (Qiagen, Manchester, UK) and was amplified (100ng) using NADH dehydrogenase I primers [4], also known as complex I (*mt9/mt11)* for mtDNA and *Ndufv1* for nuclear DNA by real-time PCR using the Power SYBR Green RT-PCR kit (Applied Biosystems, Bleisvwijk, Netherlands). The mtDNA copy number was calculated using *Ndufv1* amplification as a reference for nuclear DNA content.

**SDS-PAGE and western blotting** Antibodies used in Western (immuno-) blot analysis were the following: mouse anti-MFN1 (Abcam, ab126575, RRID:AB_11141234, 1:500), mouse anti-MFN2 (Abcam, Cambridge, UK; ab56889, RRID:AB_2142629, 1:500), goat anti-rabbit GAPDH (Cell signalling, Leiden, Netherlands; #2118s, RRID:AB_561053; 1:10 000), goat anti-mouse HRP (Abcam, Cambridge, UK; ab205719, RRID:AB_2755049, 1:5 000). Specificity of MFN1 and MFN2 antibodies was determined by MFN1/2 deletion in the islets (Fig.1B).

***In vitro* insulin secretion** Islets were isolated from mice and incubated for 1 h in Krebs-Ringer bicarbonate buffer (140 mmol/l NaCl, 3.6 mmol/l KCl, 0.5 mmol/l $NaH_2PO_4$, 2 mmol/l $NaHCO_3$ [saturated with $CO_2$], 1.5 mmol/l $CaCl_2$, 0.5 mmol/l $MgSO_4$, 10 mmol/l HEPES; pH 7.4) containing 3 mmol/l glucose. Subsequently, islets were incubated (10 islets/well) for 30-45 min in Krebs-Ringer solution with either 3 mmol/l, 10 mmol/l, 17 mmol/l glucose, 20 mmol/l glucose, 20 mmol/l KCl or 10 mmol/l glucose supplemented with 100nmol/l exendin-4 (Wuxi Apptec, Shanghai, China), the glucagon-like peptide-1 (GLP-1; Wuxi Apptect), or the glucose-dependent insulinotropic peptide (GIP; Wuxi Apptec), 10µmol/l forskolin (FSK; Sigma-Aldrich),

100μmol/l IBMX (Sigma-Aldrich), 10μmol/l H89 dihydrochloride hydrate (H89; Sigma-Aldrich) or 6μmol/l of the selective Epac activator (8-pCPT-2-O-Me-cAMP-AM, Tocris Bioscience, Bristol, UK). Secreted and total insulin content were quantified using a homogeneous time-resolved fluorescence (HTRF) insulin kit (Cisbio, Codolet, France) in a PHERAstar reader (BMG Labtech, Aylesbury, UK), following the manufacturer's guidelines or a mouse insulin ELISA kit (Alpco, Salem, USA). Data are presented as secreted insulin/insulin content or % of total insulin secreted.

**cAMP assay** Primary dispersed mouse islet cells, prepared by triturating intact islets in 0.05% trypsin/EDTA for 3 min at 37°C, were stimulated with indicated concentration of agonist or 10 μmol/l forskolin (FSK; Sigma-Aldrich) for 5 min at 37°C in serum free RPMI-1640, with 10 mmol/l glucose and 1 mmol/l IBMX (Sigma-Aldrich). At the end of each incubation, cAMP was assayed by HTRF (Cisbio cAMP Dynamic 2, Codolet, France). Data were normalised to FSK response (%).

**Single-cell fluorescence imaging** To study mitochondrial structure, cells were incubated with 100nM Mitotracker green (Thermo Fisher Scientific) in Krebs-Ringer bicarbonate buffer containing 11 mmol/l glucose for 30 min. Mitotracker green was then washed with Krebs buffer with 11 mmol/l glucose before fluorescence imaging. Single channel image stacks for Mitotracker green were recorded on a LSM780 inverted confocal microscope (Carl Zeiss, Cambridge, UK) using a ×63 1.4 numerical aperture (NA) oil objective and excitation with an argon laser (488 nm) and captured using a GaAsP (Carl Zeiss) detector. The image X,Y and Z dimensions were optimised (Nyquist settings) prior to capture images to enable post processing by deconvolution using Huygens software (Scientific Volume Imaging, Hilversum, Netherlands).

Experiments with tetramethylrhodamine ethyl ester (TMRE) were performed as previously described [5]. Traces represent mean normalised fluorescence intensity over time ($F/F_{min}$), where $F_{min}$ is the mean fluorescence recorded during the application of 3 mmol/l glucose.

Clusters of dissociated islets were transduced for 48h with an adenovirus encoding the low-$Ca^{2+}$-affinity sensor D4 addressed to the ER, Ad-RIP-D4ER (MOI: 100), as described in [6, 7]. Bleaching was corrected by fitting a simple linear equation (Microsoft Excel) to the ratio values preceding the first addition of exendin-4 (90-340s), thus generating the predicted bleaches in the absence of further additions. Corrected traces were then generated by division of the observed ratio values by those predicted by the fitted linear equation.

**Mitochondrial shape analysis** For each stack, one image at the top, middle and bottom of the islet was analysed. After background subtraction, the following parameters were measured for each cell: number of particles, perimeter and circularity of each particle and elongation (1/circularity) was calculated [8]. The average perimeter, circularity and elongation of particles was then calculated for each cell.

**Whole-islet fluorescence imaging** Mitochondrial $Ca^{2+}$ imaging of whole islets was performed after infection with adenovirus encoding the mitochondrially-targeted probe, R-GECO as previously described [5] (Addgene, Watertown, NY, USA; RRID:Addgene_46021; MOI: 100; [48 h post-isolation]) in modified Krebs-Ringer bicarbonate buffer. To examine ATP:ADP changes in response to a rise in extracellular glucose concentration, islets were infected as previously described [5] with an adenovirus bearing cDNA encoding the ATP sensor pGW1CMV-Perceval (RRID:Addgene_21737; MOI: 100; kindly provided by G. Yellen [Yale University, USA]) [9] and incubated for 48 h prior to fluorescence imaging. Imaging was performed in Krebs supplemented with 3 mmol/l, 10 mmol/l or 10 mmol/l glucose with 100nmol/l exendin-4. Cytosolic $Ca^{2+}$ imaging was performed after incubation with Cal-520 acetoxymethyl (AM; 2 µmol/l; 24 h post-isolation; Stratech, Cambridge, UK) for 45 min-1h in Krebs-Ringer bicarbonate buffer containing 3 mmol/l. Imaging was performed in Krebs supplemented with 3 mmol/l, 10 mmol/l or 17 mmol/l glucose, 10 mmol/l glucose with 100nmol/l exendin-4, 17 mmol/l glucose with 0.1 mmol/l diazoxide (Diaz; Sigma-Aldrich), or 20 mmol/l KCl with diazoxide. ER $Ca^{2+}$ storage was determined after incubation of islets with Cal-520 and 3 mmol/l glucose and imaged in Krebs buffer supplemented with 3 mmol/l or 17 mmol/l glucose, 17 mmol/l glucose with 0.1 mmol/l diazoxide or 100 μmol/l acethyl choline (Sigma-Aldrich). All images were captured at 0.5 Hz on a Zeiss Axiovert microscope equipped with a ×10 0.5 NA objective, a Hamamatsu image-EM camera coupled to a Nipkow spinning-disk head (Yokogawa CSU-10, Runcorn, UK) and illuminated at 490 nm (Cal-520, Perceval) or 580 nm (R-GECO). Data were analysed using ImageJ [10].

**TIRF fluorescence imaging** For experiments using the membrane-located zinc sensor ZIMIR [11] islets from control and dKO islets were dissociated using accutase at 37°C during 5 min and dissociated cells were left to attach on a poly-L-lysine treated glass slides for 3 hours before incubation in KREBS buffer containing 3 mmol/l glucose and ZIMIR (50 µmol/l) for 40 min. Live-imaging at the membrane was performed on a Nikon Ti (Tokyo, Japan) microscope equipped with a iLas2 TIRF (Gataca Systems, Massy, France) module and a 100x/1.49 TIRF objective at 488 nM excitation.

Acquisition rate was 3 images/second and after 3 min, KCl was added to a final concentration of 20 mmol/l.

For experiments using the fluorescent genetically-encoded and vesicle-located green marker NPY-Venus, islets were infected with an NPY-Venus adenoviral construct [12] and left for expression for 48 hours. Islets were then dissociated as described above, left to attach to glass slide and fixed using 4% PFA in PBS. TIRF imaging was performed using the same microscopy system.

**Pancreas immunohistochemistry** Isolated pancreata were fixed in 4% (vol/vol) buffered formalin and embedded in paraffin wax within 24 h of removal. Slides (5 μm) were submerged sequentially in Histoclear (Sigma-Aldrich) followed by washing in decreasing concentrations of ethanol to remove paraffin wax. Permeabilised pancreatic slices were blotted with anti-guinea pig insulin (Cell Signalling; 1:500; #4590, RRID:AB_659820) and anti-mouse glucagon (Sigma-Aldrich; 1:1 000; G2654, RRID:AB_259852) primary antibodies. Slides were visualised by subsequent incubation with Alexa Fluor 488 and 568-labelled goat anti-guinea pig and anti-mouse antibodies (Thermo Fisher Scientific; 1:1 000; #A-11073, RRID:AB_2534117 and #A-11004, RRID:AB_2534072). For examination of apoptosis, TUNEL assay was performed using a DeadEnd Fluorometric TUNEL system kit and DNase I treatment (Promega, Madison, Wisconsin, USA) according to the manufacturer's instructions. Samples were mounted on glass slides using Vectashield$^{TM}$ (Vector Laboratories, Burlington, ON, Canada) containing DAPI. Images were captured on a Zeiss AxioObserver.Z1 microscope using a ×40 Plan-Apochromat 206/0.8 M27 air objective, a CMOS ORCA Flash 4 camera (Hamamatsu, Tokyo, Japan) with a Colibri.2 LED illumination system. Fluorescent quantification was achieved using ImageJ with a purpose-designed macro (available upon request). Whole pancreas sections were used to quantitate cell mass. The number of TUNEL-positive cells of all visible islets was measured using ImageJ.

**Immunohistochemistry of Pdx1CreER pancreata** Briefly, pancreata were collected, cut into 20–30 pieces, and fixed overnight in 4% paraformaldehyde (Sigma-Aldrich) at 4°C. Tissues were then washed in PBS and frozen in OCT (Sakura Finetek, VWR International, Pennsylvania, USA) as two-blocks, with tissue-pieces randomly distributed. Blocks were sectioned at 15μm thickness, with one of every 6 sections collected onto Cryo+ (VWR International, Pennsylvania, USA) glass slides. This produces ~40 slides, with each containing 4–6 sections. Slides were used for immunofluorescence staining after antigen retrieval in pH 6 citrate buffer. Four slides from each pancreas were used for each antibody-combination. The immune-fluorescence intensities on at least 50 islet sections in each pancreas (from different slides) were assessed under an Olympus IX51 inverted fluorescence scope microscope (Olympus lifescience, Tokyo, Japan) under double-blind setting. Confocal images were taken using an Olympus FV1000 camera. Permeabilised pancreatic slices were blotted with anti-guinea pig insulin (Agilent; 1:2000; #A0564, RRID:AB_10013624), anti-rabbit glucagon (Abcam; 1:300; ab92517, RRID:AB_10561971) and anti-rabbit somatostatin (Abcam; 1: 1000; ab111912); RRID:AB_10903864) primary antibodies. Slides were visualised by subsequent incubation with Alexa Fluor 488 and 568-labelled donkey anti-rabbit and donkey anti-guinea pig antibodies (Jackson Immunoresearch; 1:1 000; #711-545-152,

RRID:AB_2313584 and #706-165-148, RRID:AB_2340460). The Vanderbilt University IACUC approved all procedures for these studies.

**Metabolomics/lipidomics** One method measuring mouse plasma molecules focused on a panel of known metabolites previously associated with diabetes, diabetes complications and metabolic dysfunction. These molecules were fully quantified using targeted ultra-high-performance liquid-chromatography coupled triple quadrupole mass spectrometry (UHPLC-QqQ-MS/MS) as described earlier [13]. The second method aimed to measure a broad array of lipid species. Lipidomic sample preparation followed the Folch procedure with minor adjustments [14]. Lipids were measured with ultra-high-performance liquid-chromatography coupled quadruple-time-of-flight mass spectrometry (UHPLC-QTOF/MS) in both positive and negative ionization mode, identification was done using MZmine (version 2.28) matching to an in-house library [15]. Peak areas were normalized to internal standards [16]. Significance was tested by Student's two-tailed t-test using GraphPad Prism 8 software.

**Measurement of oxygen consumption rate** Seahorse XF96 extracellular flux analyzer (Seahorse Bioscience, Agilent, Santa Clara, CA, USA) was used for intact mouse islets respirometry. For XF96 assays, mouse islets (~10 per well) were seeded in 1μL/well Matrigel (Corning Life Sciences, Tewksbury, MA, USA) in a poly-D-lysine (PDL)-coated (Thermo Fisher Scientific) XF96 plate and size-matched between conditions. Islets were incubated at 37 °C for 3.5min to solidify the Matrigel before the addition of 150μL/well of Seahorse assay media (XF Base Media Minimal DMEM, pH 7.4 supplemented with 3 mmol/l glucose, 200mM L-glutamine, 100mM sodium pyruvate and 0.1% FBS). Islets were incubated at 37 °C for approximately 1hr before starting the assay. 4 baseline (under 3 mmol/l glucose) OCR measurements were taken before the following substrates/compounds were injected: 20 mmol/l glucose in port A, followed by Oligomycin (Sigma-Aldrich, final concentration of 5µM) in port B, FCCP (Sigma-Aldrich, final concentration of 1µM FCCP) in port C, and Antimycin A with Rotenone (Sigma-Aldrich, final concentration of 5µM) in port D. Data are presented as pmol $O_2$/min/islets.

**Electron microscopy (EM)** For conventional EM, islets were chemically fixed in 2% paraformaldehyde (EM grade, TAAB Laboratories Equipment, Berks, UK), 2% glutaraldehyde and 3 mM $CaCl_2$ in 0.1 M cacodylate buffer (Sigma-Aldrich) for 2 h at room temperature then left overnight at 4°C in fresh fixative solution, osmicated, enrobed in agarose plugs, dehydrated in ethanol and embedded on Epon (TAAB Laboratories Equipment). Epon was polymerised overnight at 60°C. Ultrathin 70 nm sections were cut with a diamond knife (DiATOME, Nidau, Switzerland) in a Ultracut UCT ultramicrotome (Leica Biosystem Technologies, Amsterdam, Netherlans) before examination on a Tecnai T12 TEM (FEI, Lausanne, Switzerland). Images were acquired in a charge-coupled device camera (Quantum Design, Surrey, UK), and processed in ImageJ.

***In vivo* $Ca^{2+}$ imaging of AAV8-INS-GCaMP6s infected endogenous pancreatic islets** All mice were i.p. injected with tamoxifen daily at 15 weeks old. Endogenous pancreatic islets of control and β*Mfn1/2*-KO mice (16 weeks old) were then infected with AAV8-INS-GCaMP6s [17] viral particles ($1*10^{14}$ or $10.1*10^{13}$ genome copies/ml) via i.p. injection (50µl/mouse), 24-28 days prior to terminal intravital imaging. On the day of imaging, mice were anesthetized with 2-4% inhaled isoflurane, the pancreas was then externalised and placed on a coverslip on a TCS SP8 DIVE (Leica Biosystem Technologies) multiphoton platform equipped with heating pads and an objective warmer. Heated blankets were used at all times and temperature was constantly monitored using a rodent rectal temperature probe throughout the imaging session. A ×40 1.1 NA water immersion objective was used during imaging. Islets were identified using green fluorescence from the GCaMP6s biosensor. Dyes suspended in saline, including Hoechst 33342 (Thermo Fisher Scientific: H3570; 70μg), tetramethylrhodamine methyl ester (TMRM, Thermo Fisher Scientific; T668; 3μg) and Alexa Fluor 647 NHS Ester (Thermo Fisher Scientific; A20006) conjugated to Rat serum Albumin (Albumin-647; 500μg) were injected retro-orbitally to label nuclei, mitochondria and vasculature, respectively. *In vivo* $Ca^{2+}$ imaging was performed by simultaneously exciting GCaMP6s at both 820 nm and 940 nm to capture non-excited and excited states of $Ca^{2+}$ activity in the beta cells. Fluorescent emission was detected by an external HyD detector (Leica Biosystem Technologies) through a 500-550 nm bandpass filter. Hoechst, TMRM and the dual excitable Albumin-647 were excited at 820 nm and fluorescence emission was detected by either external PMT (Leica Biosystem Technologies) or HyD detectors through 400-470 nm, 570-630 nm and 650-750 nm bandpass filters, respectively. Baseline imaging of islets were recorded 10 min prior i.p. injection of glucose (1g/kg in sterile saline). The same islets were imaged for ~20 min to monitor $Ca^{2+}$ oscillations post glucose injection. Baseline and post-imaging glucose measurements were obtained using an AlphaTRAK 2 (Zoetis Services, Leatherhead, UK) glucometer. All images were collected at a scan speed of 600 Hz with a line average of 2, and a 512 x 512 frame size, with a frame rate of 0.576/s. At the end of each imaging session, animals were perfused with 4% (vol/vol) PFA and tissues were paraffin-embedded for further analysis. All *in vivo* imaging experiments were performed with approval and oversight from the Indiana University Institutional Animal Care and Use Committee (IACUC).

Fluorescent traces were calibrated to calculate fluorescence spikes fold change above the baseline on three cells per condition using ImageJ. Then the AUC was determined, with a threshold at 0.15 to subtract background noise.

**Connectivity analysis**

**Pearson (*r*)-based connectivity and correlation analyses** Correlation analyses in an imaged islet were performed between beta cell pairs and their extracted fluorescent $Ca^{2+}$ traces over time in MATLAB (MathWorks, Cambridge, UK) using a custom-made script (available on request) as previously described [18]. A noise reduction function (effectively a rolling average) was applied to smooth noisy data and minimize the effects of outliers. The data window size used to calculate the moving averages was set to 5% of the total data points collected during each capture. The correlation coefficient *r* between all possible (smoothed) beta cell pair combinations (excluding the autocorrelation) was assessed using Pearson's function. The Cartesian co-ordinates of the imaged cells were then incorporated in the construction of connectivity line maps. Cell pairs were connected with a straight line, the colour of which represented the correlation strength and was assigned to a colour-coded light-dark ramp (R=0.1-0.25 [blue], 0.26-0.5 [green], R=0.51-0.75 [yellow], R=0.76-1.0 [red]). Cells with the highest number of possible cell pair combinations are shown in red. Data are also displayed as heatmap matrices, indicating all possible beta cell pair connections on each axis (min. = 0; max. = 1). The positive r values (excluding the auto-correlated cells) and the percentage of cells that were connected to one another were averaged and compared between groups.

**Monte Carlo-based signal binarisation and data shuffling for identification of highly connected cells** Data were analysed using approaches similar to those previously described [19, 20]. $Ca^{2+}$ signals were denoised by subjecting the signal the Huang-Hilbert type (HHT) empirical mode decomposition (EMD). The signals were decomposed into their intrinsic mode functions (IMFs) in MATLAB (MathWorks) [21]. The residual and the first IMF with the high-frequency components were then rejected to remove random noise.

The Hilbert-Huang Transform was then performed to retrieve the instantaneous frequencies [22-24] of the other IMFs to reconstruct the new signal using

$$X(t) = Re \sum_{j=1}^{N} a_j(t) e^{i \int \omega_j(t) dt}$$

where $a_j(t)$ = amplitude, $\omega_j(t)$ = frequency of the *i*th IMF component to retrieve a baseline trend and to account for any photobleaching or movement artefacts. A 20% threshold was imposed to minimise false positives from any residual fluctuations in baseline fluorescence.

Cell signals with deflection above the de-trended baseline were represented as '1' and inactivity represented as '0', thus binarising the signal at each time point. The coactivity of every cell pair was then measured as:

$$C_{ij} = \frac{T_{ij}}{\sqrt{T_i T_j}}$$

where $T_{ij}$ = total coactivity time, $T_i$ and $T_j$ = total activity time for two cells.

The significance at $p<0.001$ of each coactivity measured against chance was assessed by subjecting the activity events of each cell to a Monte Carlo simulation [25, 26] with 10,000 iterations.

Synchronised $Ca^{2+}$-spiking behaviour was assessed by calculating the percentage of coactivity using the binarised cell activity dataset. A topographic representation of the connectivity was plotted in MATLAB (MathWorks) with the edge colours representing the strength of the coactivity between any two cells.

An 80% threshold was imposed to determine the probability of the data, which was then plotted as a function of the number of connections for each cell to determine if the dataset obeyed a power-law relationship [27].

**RNA-Seq data analysis** Processing and differential expression analysis of RNA-Seq data from high fat high sugar (HFHS, D12331, Research Diets) and regular chow (RC) fed mice was performed as previously described [28]. Briefly, differentially expressed genes were computed for the HFHS vs RC comparisons at 4 time-points for each of six mouse strains ((C57Bl/6J, DBA/2J, BALB/cJ, A/J, AKR/J, 129S2/SvPas) using the *Limma* package in R and p-values were adjusted for multiple comparisons using the Benjamini Hochberg procedure [29].

### Supplemental Figure legends

**Supplemental Fig.1 Pdx1CreER activation has no detectable effect on glycaemia both *in vivo* and *in vitro* and on key beta-cell gene expression and islet morphology.** (A) Glucose tolerance measured by IPGTT (2 g/kg body weight) in WT and Pdx1CreER mice (*n*=6 mice per genotype) at 8 weeks of age. (B) Insulin secretion measured during serial incubations in batches in 3 or 20 mmol/l glucose (*n*=6 mice per genotype in three independent experiments) at 8 weeks of age. (C) (a,b) MafA (red) and insulin (green) expression levels in WT and Pdx1CreER islets. (c,d) Typical distribution of beta- (insulin, red), alpha- (glucagon, green), and delta-cells (Somatostatin, SS, green) in 8-week old islet sections (*n*= 50 islets, 3 male mice per genotype). Note that both alpha and delta cells were unidentifiable as these were stained in green. Scale bar: 20 μm. Data are presented as mean±SEM. Data assessed by two-way ANOVA test and Sidak's multiple comparisons test.

**Supplemental Fig.2 Body weight loss, insulin resistance and increased β-ketone production is observed in β*Mfn1/2* dKO mice.** (A) Measured body weight in control and β*Mfn1/2* dKO mice (*n*=3-6 mice per genotype) at 14 weeks of age. (B) Glucose tolerance measured by IPGTT (1 g/kg body weight) in 20-week-old mice and (C) the corresponding AUC were assessed in β*Mfn1/2* dKO and control mice (*n*=8 mice per genotype, in 2 independent experiments). (D) Challenging β*Mfn1/2* dKO mice with a 0.75 U/kg body weight insulin injection as compared with control mice at 14 weeks of age. Data normalised to baseline (%). (E) Corresponding AUC for (D) is also shown (*n*=6 mice per genotype). (F) Glucose and (G) β-ketone bodies measured before or after an overnight (16h) fasting in 14-week control and dKO mice. (H) Plasma insulin levels were quantified under fed and fasted conditions in 14-week dKO and control mice (*n*=6 mice per genotype). Data are presented as mean±SEM. *$p<0.05$; **$p<0.01$; ***$p<0.001$; ****$p<0.0001$ as indicated, or at the time points indicated analysed by unpaired two-tailed Student's t-test and Mann–Whitney correction or two-way ANOVA test and Sidak's multiple comparisons test. Experiments were performed in 14 or 20-week-old male mice as stated accordingly.

**Suppl. Fig.3 Heatmap of differential gene expression between β*Mfn1/2* dKO and control islet mRNA**. Changes in key beta or alpha cell genes, disallowed genes, mitochondrial, ER stress or mito/autophagy genes were assessed by qRT-PCR in control and dKO islets according to the colour coded median values from 0 to 1, white to dark blue respectively (*n*=3-4 mice per genotype; experiment performed in duplicate). Expression values for each gene were normalised to β-*actin*. *p<0.05; **p<0.01, assessed by two-way ANOVA test and Sidak's multiple comparisons test. Experiments were performed in 14-week-old male mice.

**Supplemental Fig.4 Mfn1/2 deletion impairs beta cell function *in vivo*. Representative *in vivo* images of GCaMP6s labelled islets and TMRM stained mitochondria surrounded by their vasculature in control and β*Mfn1/2* dKO mice.** (A) Representative traces depicting fluorescence intensity of cytosolic $Ca^{2+}$ (GCaMP6s) and mitochondrial TMRM signals in control and (B) dKO animals; scale bar: 45 μm; (*n*=1 animal per genotype). Parallel and anti-parallel fluorescent signals are shown within selected areas. (C) AUC of fold change measurements above baseline for each GCaMP6s and TMRM traces measured (*n*=3 total responding cells). Green, GCaMP6s; red, TMRM signals. Islets were imaged in 20 week-old mice. Data are presented as mean±SEM. *p<0.05, assessed by two-way ANOVA test and Sidak's multiple comparisons test.

**Supplemental Fig.5 Impact of *Mfn1/2* deletion on intercellular connectivity.** (A) Representative cartesian maps of islets with colour coded lines connecting cells according to the strength of Pearson analysis (colour coded *r* values from 0 to 1, blue to red respectively) under 3mmol/L (3G), 17mmol/L (17G) glucose or 20mmol/L KCl; scale bars: 40 μm.(B) Representative heatmaps depicting connectivity strength (*r*) of all cell pairs according to the colour coded *r* values from 0 to 1, blue to yellow respectively.(C) Percentage of correlated cell pairs at 3G, 17G or KCl (*n*=17-26 islets, 4 mice per genotype).(D) *r* values between beta cells in response to glucose or KCl (*n*=4 mice per genotype).(E) qRT-PCR quantification of *Cx36* expression relative to β-*actin* (*n*=3–4 mice per genotype in two independent experiments). Data are presented as mean±SEM. *p<0.05, assessed by unpaired two-tailed Student's t-test and Mann–Whitney correction or two-way ANOVA test and Sidak's multiple comparisons test. Analysis and experiments were performed on data collected from 14-week-old male mice.

**Supplemental Fig.6 Mapping of beta cell population dynamics.** (A) Representative cartesian maps of islets stained with Cal-520 with colour coded lines connecting cells according to the strength of coactivation (colour coded *r* values from 0 to 1, blue to red respectively) under 3mmol/L (3G), 17mmol/L (17G) glucose or 20mmol/L KCl (B) Log-log graph of the beta cell-beta cell connectivity distribution at 17 mmol/L glucose. control (*n*=17 islets) and dKO (*n*=26 islets) beta cells display obedience to a power-law distribution whereby few beta cells host 60% to 100% of the connections to the rest of the beta cell population (*n*=4 animals per genotype; experiment performed in triplicate). Analysis was performed on data collected from 14-week-old male mice.

**Supplemental Fig.7 Impaired insulin secretion observed in Clec16a$^{\Delta panc}$ islets can be rescued by GLP-1R agonists *in vitro.*** (A) Insulin secretion measured in control (Pdx1-Cre) and Clec16a$^{\Delta panc}$ mice in 3 mmol/l glucose (3G), 17 mmol/l glucose (17G), or 10 nmol/l exendin-4 (ex4) (*n*=4 mice per genotype). (B) Glucose tolerance measured by IPGTT (1.5 g/kg body weight) in 8-week-old male Pdx1-Cre and Clec16a$^{\Delta panc}$ mice or OGTT (1.5 g/kg body weight) in 9–10-week-old animals. (C) The corresponding AUC is shown in (B) (*n*=4-5 mice per genotype). (*p<0.05,**p<0.01, control OGTT vs Clec16a$^{\Delta panc}$; #p<0.05, ##p<0.01, control IPGTT vs Clec16a$^{\Delta panc}$). Data are presented as mean±SEM and assessed by two-way ANOVA test and Sidak's multiple comparisons test.

**Supplemental Fig.8 Insulin granule density is increased in β*Mfn1/2* dKO beta cells.** (A) Confocal images of NPY-Venus fluorescence in dissociated fixed pancreatic beta cells isolated from control and dKO mice. Scale bar: 10 µm.(B) Effect of KCl on exocytosis as reported with NPY-Venus in pancreatic beta cells. Traces represent mean normalised fluorescence intensity over time ($F/F_{min}$). (C) Confocal images of ZIMIR fluorescence imaging in dissociated pancreatic beta cells isolated from control and dKO mice. Scale bar: 10 µm.(D) Representative time courses of ZIMIR signal fold change above baseline ($F/F_{min}$) upon KCl-stimulated insulin/$Zn^{2+}$ release and (E) fold change of peaks in dissociated control and dKO cells. (*n*=19 cells from 3 control mice; *n*=12 cells from 3 β*Mfn1/2* dKO mice). Data are presented as mean±SEM. *p<0.05,**p<0.01; assessed by unpaired two-tailed Student's t-test and Mann–Whitney correction or two-way ANOVA test and Sidak's multiple comparisons test. Experiments were performed in 14-week-old male mice.

**Supplemental Fig.9 Volcano plots showing alterations in metabolites and lipids from plasma samples of control and β*Mfn1/2* dKO mice.** (A) Volcano plot summarising both fold-change and t-test criteria for all metabolites. Results are summarised in a scatter-plot of the negative $\log_{10}$-transformed p values from the t-test plotted against the $\log_2$ fold change. Negative values indicate downregulated metabolites in dKO mice, while positive values reflect upregulated metabolites. Metabolites with statistically significant differential levels according to the t-test lie above a horizontal threshold line (red dots). Metabolites with large fold-change values lie far from the vertical threshold line at $\log_2$ fold change = 0, indicating whether the metabolite is up or downregulated. The list of analysed metabolites with their abbreviations is presented in ESM table 3. (B) Lipids that were found downregulated in dKO mice with statistically significant differential levels according to the t-test are presented above a horizontal threshold line. Plasma samples were isolated from *n*=3 animals per genotype. The most significantly downregulated lipids are annotated. SM, sphingomyelins; CER, ceramide; CE, cholesterol esters; DG, di(acyl/alkyl)glycerols; FA, fatty acids; TG, tri(acyl/alkyl)glycerols; LPC, lysophosphatidylcholines; PC, phosphatidylcholines;LPE,lysophosphatidylethanolamines;PE,phosphatidylethanolamines; PG, phosphatidylglycerols; PI, phosphatidylinositols; PS, phosphatidyserines. Experiments were performed in 14-week-old male mice.

**Supplemental Fig.10 Boxplots showing differences between HFHS (yellow) and RC (green) diet in 6 mouse strains over time for *Mfn1* (A) and *Mfn2* (B) genes.** The bottom and top of the boxes represent the first and third quartiles, with the horizontal line representing the median. The upper whiskers represent the third quartile plus 1.5x IQR (interquartile range); the lower whiskers represent the first quartile minus 1.5x IQR. Outlier points beyond this range are indicated above or below the whiskers. Statistically significant comparisons following false discovery rate (FDR) correction (FDR <= 0.05) are indicated by a double asterisk. Marginally significant comparisons (raw p value <= 0.05) are indicated by a single asterisk.

### Electronic Supplemental Tables

#### ESM Table 1

| Genes | Forward primer (5’→3’) | Reverse Primer (5→3’) |
|---|---|---|
| *Mfn1* | TGGTAATCTTTAGCGGTGCTC | GGAGGACTTTATCCCACAGC |
| *Mfn2* | TTTGGAAGTAGGCAGTCTCCA | CAGGCAGCACTGAAAAGAGA |
| *Pdx1-Cre*$^{ERT2}$ | CAGGCGTTTTCTGAGCATACC | CCGGTTATTCAACTTGCACCAT |

Sequence of primers used for genotyping *Mfn1* and *Mfn2* flox.

### ESM Table 2

| Genes | Forward primer (5’→3’) | Reverse Primer (5→3’) |
|---|---|---|
| *Mfn1* | GCATTTTTTGGCAGGACAAGTAG | GGAGGACTTTATCCCACAGCAT |
| *Mfn2* | AGAAGAGTGTCAAGACTGTGAACCA | GCTGCCTGCATGCAACTG |
| *Drp1* | TCAGATCGTCGTAGTGGGAA | TCTTCTGGTGAAACGTGGAC |
| *Opa1* | ATACTGGGATCTGCTGTTGG | AAGTCAGGCACAATCCACTT |
| *Fis1* | AAGTATGTGCGAGGGCTGT | TGCCTACCAGTCCATCTTTC |
| *Ins2* | TGGCTTCTTCTACACACCCATGTCCC | ACTGATCTACAATGCCACGCTTCTGCT |
| *Slc2a2* | GCAACTGGGTCTGCAATTTTG | CAAGGAAGTCCGCAATGTACTG |
| *Gck* | TGGTGGATGAGAGCTCAGTGAA | CATGTACTTTCCGCCAATGATC |
| *Pdx1* | CCAAAGCTCACGCGTGGA | TGTTTTCCTCGGGTTCCG |
| *Nkx6.1* | GCCTGTACCCCCCATCAAG | GTGGGTCTGGTGTGTTTTCTCTT |
| *Pax6* | GCACATGCAAACACACATGAAC | GGTGAAATGAGTCCTGTTGAAGTG |
| *Ucn3* | GCTGTGCCCCTCGACCT | TGGGCATCAGCATCGCT |
| *Glp1r* | CCCTGGGCCAGTAGTGTG | GCAGGCTGGAGTTGTCCTTA |
| *Nkx2.2* | CCTCCCCGAGTGGCAGAT | GAGTTCTATCCTCTCCAAAAGTTCAAA |
| *Gcg* | TCACAGGGCACATTCACCAG | CATCATGACGTTTGGCAATGTT |
| *Arx* | TCCGGATACCCCACTTAGCTT | GACGCCCCTTTCCTTTAAGTG |
| *Mafa* | CTTCAGCAAGGAGGAGGTCATC | CGTAGCCGCGGTTCTTGA |
| *Mafb* | TGAATTTGCTGGCACTGCTG | AAGCACCATGCGGTTCATACA |
| *Vdac1* | GCTAAGGATGACTCGGCTTTAAGG | AGGTTAAGTGATGGGCTAGGATGG |
| *Vdac2* | TCACTGTTGGCTGGTTCCTAGTTG | AAGACCTCGTGGATTATGCTAGGG |
| *Vdac3* | CACTTGTCCCTGGAAATGAAGAG | CATGACACTACGTTGTTGCTGAGG |
| *Letm1* | TCCTGCGTTTCCAGCTCACCAT | GTCTTCTGTGACACCGAGAGCT |
| *Slc8a1* | CCGTGACTGCCGTTGTGTT | GCCTATAGACGCATCTGCATACTG |
| *Trpm5* | CCAGCATAAGCGACAACATCT | GAGCATACAGTAGTTGGCCTG |
| *Beclin-1* | TGGAAGGGTCTAAGACGT | GGCTGTGGTAAGTAATGGA |
| *Lc3* | CACTGCTCTGTCTTGTGTAGGTTG | TCGTTGTGCCTTTATTAGTGCATC |
| *Bnip3* | TTCCACTAGCACCTTCTGATGA | GAACACCGCATTTACAGAACAA |
| *p62* | CCCAGTGTCTTGGCATTCTT | AGGGAAAGCAGAGGAAGCTC |

| | | |
|---|---|---|
| *GabarapL* | CATCGTGGAGAAGGCTCCTA | ATACAGCTGGCCCATGGTAG |
| *CathepsinL* | GTGGACTGTTCTCACGCTCAAG | TCCGTCCTTCGCTTCATAGG |
| *Pink1* | TGAGGAGCAGACTCCCAGTT | AGTCCCACTCCACAAGGATG |
| *Parkin* | TGGAAAGCTCCGAGTTCAGT | CCTTGTCTGAGGTTGGGTGT |
| *Atf4* | GCAGTGTTGCTGTAACGGACA | CGCTGTTCAGGAAGCTCATCT |
| *Atf6a* | GACTCACCCATCCGAGTTGTG | CTCCCAGTCTTCATCTGGTCC |
| *Bip* | AGGACAAGAAGGAGGATGTGGG | ACCGAAGGGTCATTCCAAGTG |
| *Chop2* | CCACCACACCTGAAAGCAGAA | AGGTGAAAGGCAGGGACTCA |
| *Xbp1* | TGGCCGGGTCTGCTGAGTCCG | GTCCATGGGAAGATGTTCTGG |
| *Xbp1s* | CTGAGTCCGAA TCAGGTGCAG | GTCCATGGGAAGATGTT CTGG |
| *Cx36* | CAGCAGCACTCCACTATGATTG | GTACACCGTCTCCCCTACAA |
| *Ldha* | ATGAAGGACTTGGCGGATGA | ATCTCGCCCTTGAGTTTGTCTT |
| *Slc16a1* | GCTTGGTGACCATTGTGGAAT | CCCAGTACGTGTATTTGTAGTCTCCAT |
| *Pdgfra* | GACCCTGTTCCAGAGGAGGAA | TTCCGAAGTCTGTGAGCTGTGT |
| *Aldh1a3* | GGGCCTCAGATCGACCAAAA | CTAGCTTGGCCCCTTCCTTC |
| *Hsd11b* | GGAGCCGCACTTATCTGA | TGCCATTTCTCTTCCAATC |
| *Mt9/mt11* | GAGCATCTTATCCACGCTTCC | GGTGGTACTCCCGCTGTAAA |
| *Ndufv1* | CTTCCCCACTGGCCTCAAG | CCAAAACCCAGTGATCCAGC |
| *β-actin* | CGAGTCGCGTCCACCC | CATCCATGGCGAACTGGTG |

List of primers used for qRT-PCR.

**ESM Table 3**

| Metabolites | Abbreviations | control mean | dKO mean | $log_2$ (fold change) | Student t-test (p value) |
|---|---|---|---|---|---|
| AADA | Aminoadipic acid | 1909.1 | 699.12 | -1.4493 | 0.1347 |
| a(R)-OHB/a(S)-OHB | Alpha-hydroxybutyric acid | 2000.2 | 2105.2 | 0.0738 | 0.9064 |
| ADMA/SDMA | (A)symmetric dimethylarginine | 151.88 | 152.32 | 0.0042 | 0.9968 |
| Ala | Alanine | 50606 | 35866 | -0.4967 | 0.3161 |
| β-OHB | β-hydroxybutyric acid | 3990.9 | 5578.9 | 0.4833 | 0.4692 |
| CA | Cholic acid | 1181.3 | 4177.4 | 1.8222 | 0.1482 |
| CDCA | Chenodeoxycholic acid | 327.1 | 348.92 | 0.0931 | 0.2608 |
| Cit | Citrulline | 6926.6 | 6273.8 | -0.1428 | 0.5838 |
| DCA | Deoxycholic acid | 216.51 | 252.52 | 0.222 | 0.266 |
| GBB | Gamma-butyrobetaine | 801.64 | 760.72 | -0.0756 | 0.6991 |
| GCA | Glycocholic acid | 547.92 | 655.65 | 0.259 | 0.0166 |
| Gln | Glutamine | 97592 | 93082 | -0.0683 | 0.7184 |
| Glu | Glutamic acid | 21063 | 17968 | -0.2293 | 0.5822 |
| Gly | Glycine | 20756 | 16462 | -0.3344 | 0.1185 |
| GUDCA | Glycoursodeoxycholic acid | 492.84 | 492.78 | -0.0002 | 0.8557 |
| HCit | Homocitrulline | 991.28 | 979.06 | -0.0179 | 0.3277 |
| Ile | Isoleucine | 14195 | 19491 | 0.4575 | 0.0196 |
| IndS | Indoxyl sulfate | 8683.7 | 6507.5 | -0.4162 | 0.119 |
| Kynu | Kynurenine | 714.87 | 729.99 | 0.0302 | 0.4022 |
| Leu | Leucine | 17020 | 21708 | 0.351 | 0.029 |
| N-MNA | N-methylnicotineamide | 547.54 | 547.58 | 1E-04 | 0.3752 |
| Phe | Phenylalanine | 11509 | 11749 | 0.0298 | 0.8802 |
| Taurine | Taurine | 128076 | 100512 | -0.3496 | 0.4514 |
| TCA | Taurocholic acid | 30302 | 110668 | 1.8687 | 0.0522 |
| TDCA/TCDCA | Tauro(cheno)deoxycholic acid | 723.01 | 1904.5 | 1.3973 | 0.0056 |
| Trp | Tryptophan | 8152.9 | 8051.9 | -0.018 | 0.9343 |
| TUDCA | Tauroursodeoxycholic acid | 548.74 | 640.49 | 0.223 | 0.3052 |
| Tyr | Tyrosine | 12364 | 12391 | 0.0031 | 0.9929 |
| UDCA | Ursodeoxycholic aicd | 418.55 | 437.05 | 0.0624 | 0.2671 |

Metabolite differences found in plasma samples of control vs dKO mice according to metabolic class and both fold-change and t-test criteria.

### ESM Videos

#### ESM Video 1

Intravital fluorescence imaging of cytosolic $Ca^{2+}$ (GCaMP6s, green), mitochondrial TMRM (red) signals and vessels (Albumin 647, white) in control animals before and after (25 sec.) glucose injection.

#### ESM Video 2

Intravital fluorescence imaging of cytosolic $Ca^{2+}$ (GCaMP6s, green), mitochondrial TMRM (red) signals and vessels (Albumin 647, white) in β*Mfn1/2* dKO animals before and after (25 sec.) glucose injection.

#### ESM Video 3

Fluorescence imaging of cytosolic $Ca^{2+}$ oscillations using Cal-520 in control (left) and β*Mfn1/2* dKO (right) whole islets in response to 3G, 3 mmol/l glucose, 17 mmol/l glucose (17G; with or without diazoxide [diaz]) or 20 mmol/l KCl with diaz. Scale bars: 50µm.

#### ESM Video 4

Fluorescence imaging of mitochondrial $Ca^{2+}$ oscillations using R-GECO in control (left) and β*Mfn1/2* dKO (right) whole islets in response to 3G, 3 mmol/l glucose, 17 mmol/l glucose (17G; with or without diazoxide [diaz]) or 20 mmol/l KCl with diaz. Scale bars: 50µm.

#### ESM Video 5

Changes in $[Ca^{2+}]_{ER}$ were measured by fluorescence imaging of cytosolic $Ca^{2+}$ oscillations using Cal-520 in control (left) and β*Mfn1/2* dKO (right) whole islets in response to 3G, 3 mmol/l glucose, 17 mmol/l glucose (17G; with or without diazoxide [diaz] or Acethylcholine [Ach]) or 20 mmol/l KCl with diaz. Scale bars: 50µm.

#### ESM Video 6

Fluorescence imaging of cytosolic $Ca^{2+}$ oscillations using Cal-520 in control (left) and β*Mfn1/2* dKO (right) whole islets in response to 3G, 3 mmol/l glucose, 10 mmol/l glucose (10G; with or without Exendin-4 [ex4]) or 20 mmol/l KCl. Scale bars: 50µm.

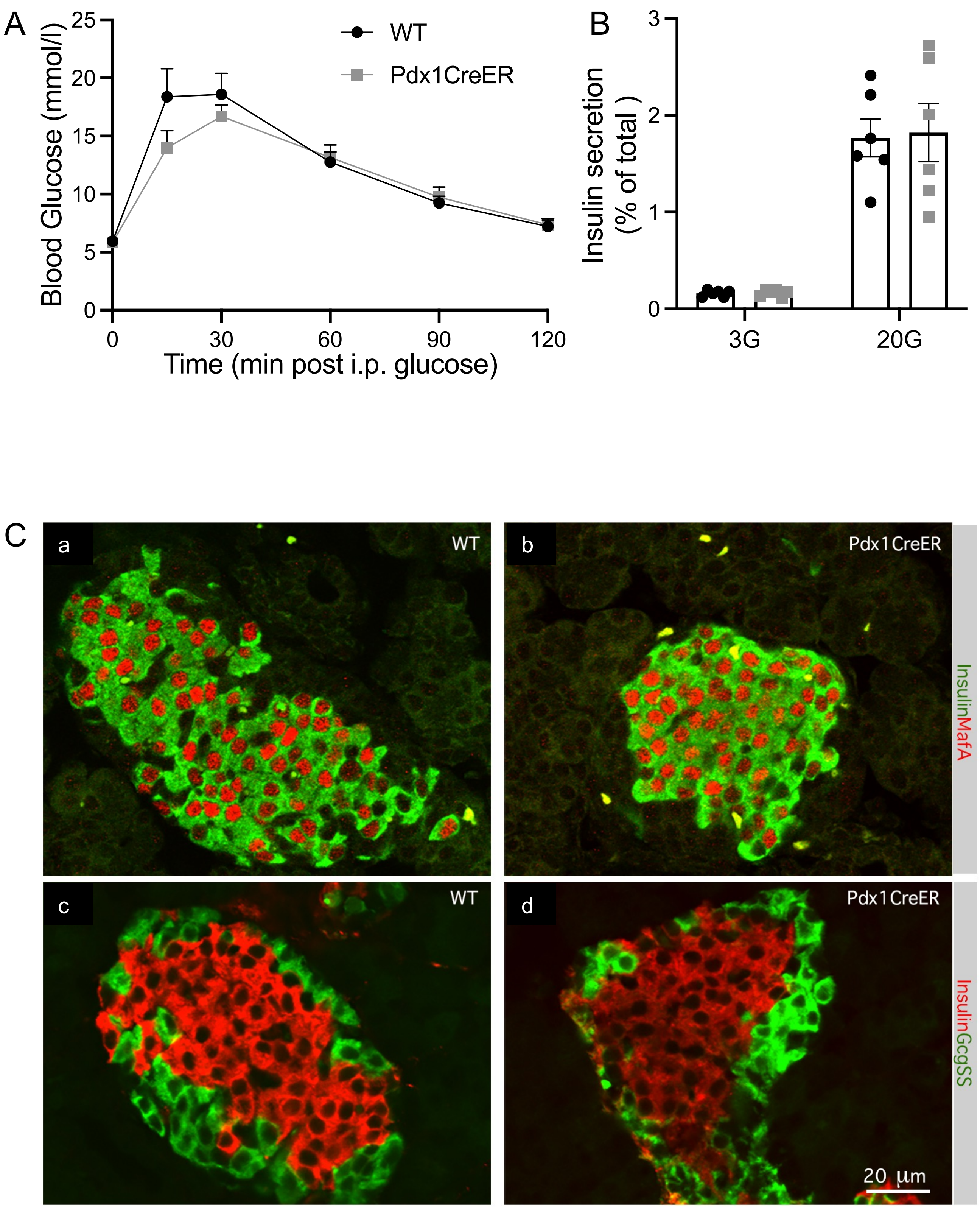
A
Blood Glucose (mmol/l)
25
20
15
10
5
0
0
30
60
90
120
Time (min post i.p. glucose)
WT
Pdx1CreER
B
Insulin secretion
(% of total )
3
2
1
0
3G
20G
C
a
WT
b
Pdx1CreER
InsulinMafA
c
WT
d
Pdx1CreER
InsulinGcgSS
20 µm

Suppl. Fig.1

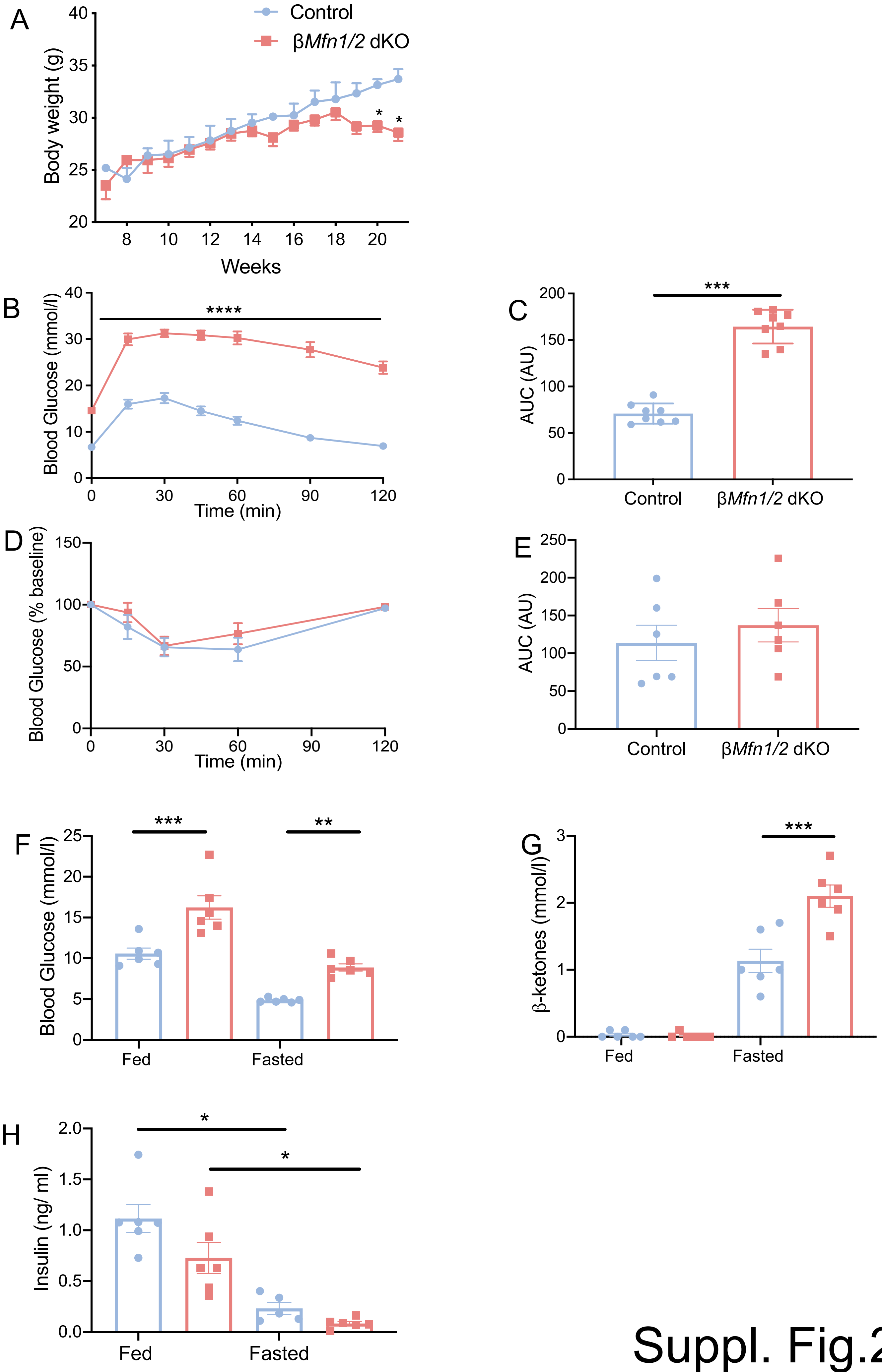
A
Control
βMfn1/2 dKO
Body weight (g)
Weeks
B
Blood Glucose (mmol/l)
Time (min)
****
C
AUC (AU)
***
Control
βMfn1/2 dKO
D
Blood Glucose (% baseline)
Time (min)
E
AUC (AU)
Control
βMfn1/2 dKO
F
Blood Glucose (mmol/l)
***
**
Fed
Fasted
G
β-ketones (mmol/l)
***
Fed
Fasted
H
Insulin (ng/ ml)
*
*
Fed
Fasted

Suppl. Fig.2

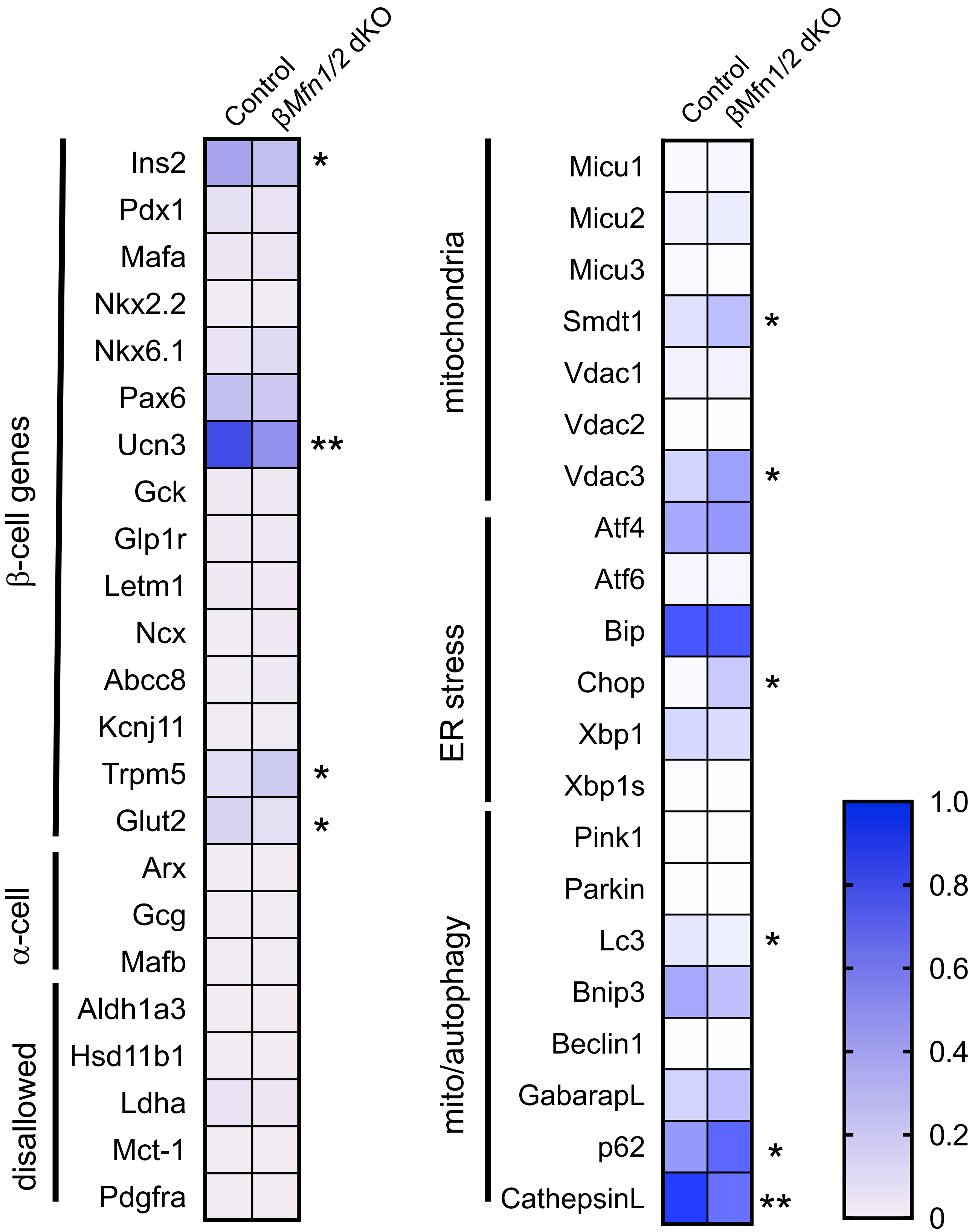
Control
βMfn1/2 dKO
β-cell genes
Ins2 *
Pdx1
Mafa
Nkx2.2
Nkx6.1
Pax6
Ucn3 **
Gck
Glp1r
Letm1
Ncx
Abcc8
Kcnj11
Trpm5 *
Glut2 *
α-cell
Arx
Gcg
Mafb
disallowed
Aldh1a3
Hsd11b1
Ldha
Mct-1
Pdgfra
Control
βMfn1/2 dKO
mitochondria
Micu1
Micu2
Micu3
Smdt1 *
Vdac1
Vdac2
Vdac3 *
ER stress
Atf4
Atf6
Bip
Chop *
Xbp1
Xbp1s
mito/autophagy
Pink1
Parkin
Lc3 *
Bnip3
Beclin1
GabarapL
p62 *
CathepsinL **
1.0
0.8
0.6
0.4
0.2
0

Suppl. Fig.3

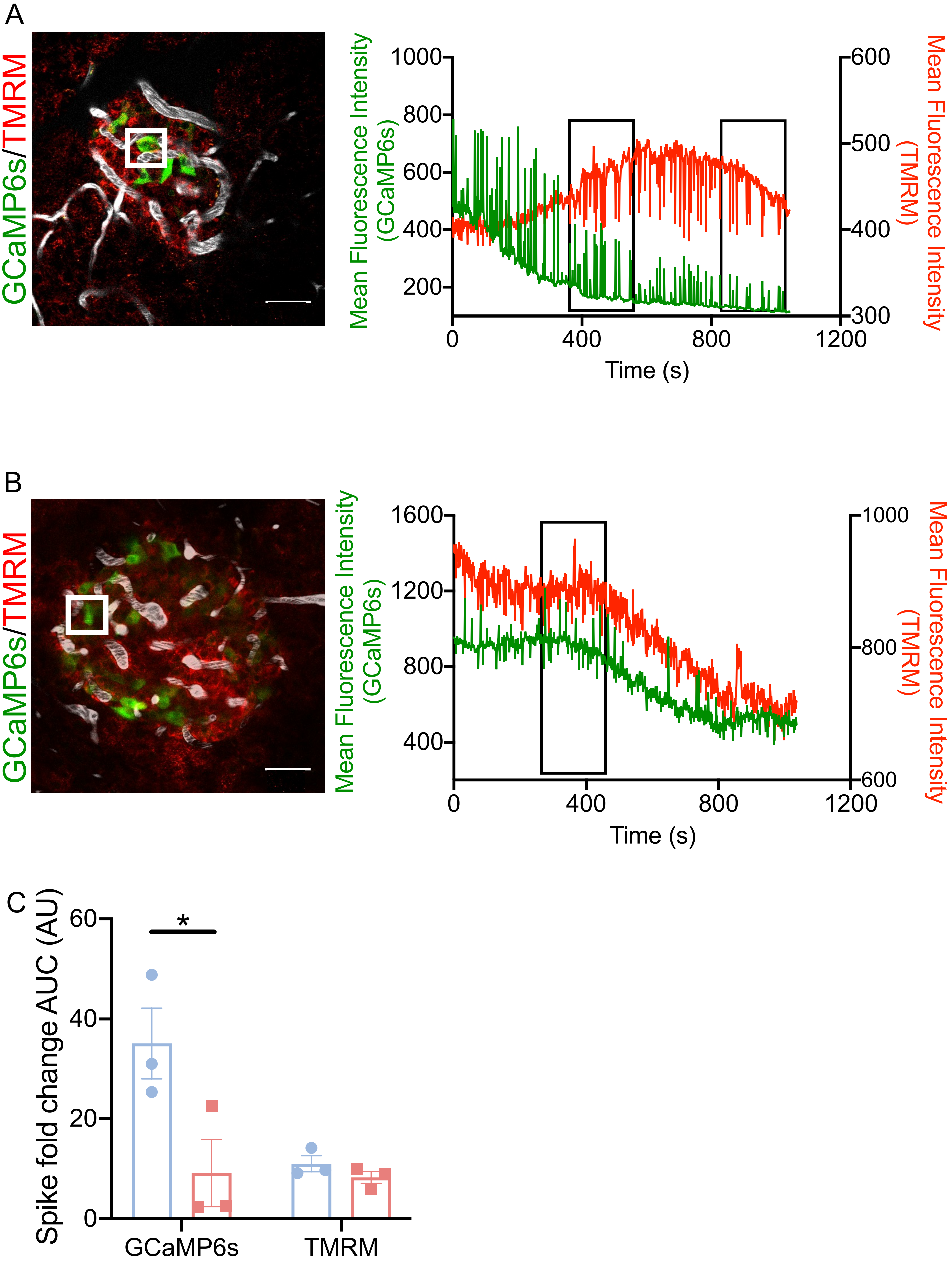

A
GCaMP6s/TMRM
Mean Fluorescence Intensity (GCaMP6s)
Mean Fluorescence Intensity (TMRM)
1000
800
600
400
200
600
500
400
300
0
400
800
1200
Time (s)
B
GCaMP6s/TMRM
Mean Fluorescence Intensity (GCaMP6s)
Mean Fluorescence Intensity (TMRM)
1600
1200
800
400
1000
800
600
0
400
800
1200
Time (s)
C
Spike fold change AUC (AU)
60
40
20
0
*
GCaMP6s
TMRM


Suppl. Fig.4

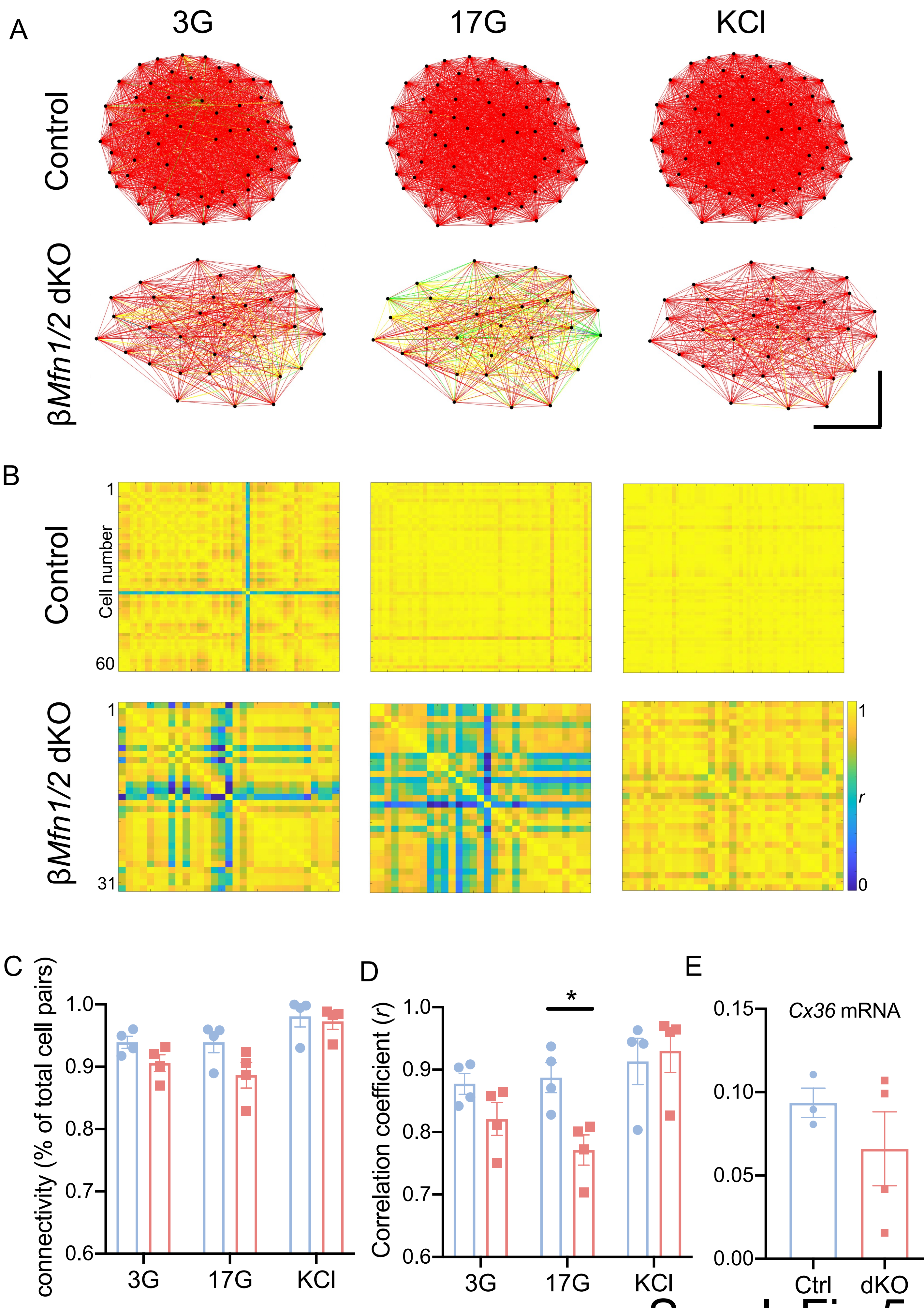

A
3G
17G
KCl
Control
βMfn1/2 dKO
B
Control
Cell number
1
60
βMfn1/2 dKO
1
31
1
r
0
C
connectivity (% of total cell pairs)
1.0
0.9
0.8
0.7
0.6
3G
17G
KCl
D
Correlation coefficient (r)
1.0
0.9
0.8
0.7
0.6
*
3G
17G
KCl
E
Cx36 mRNA
0.15
0.10
0.05
0.00
Ctrl
dKO


Suppl. Fig.5

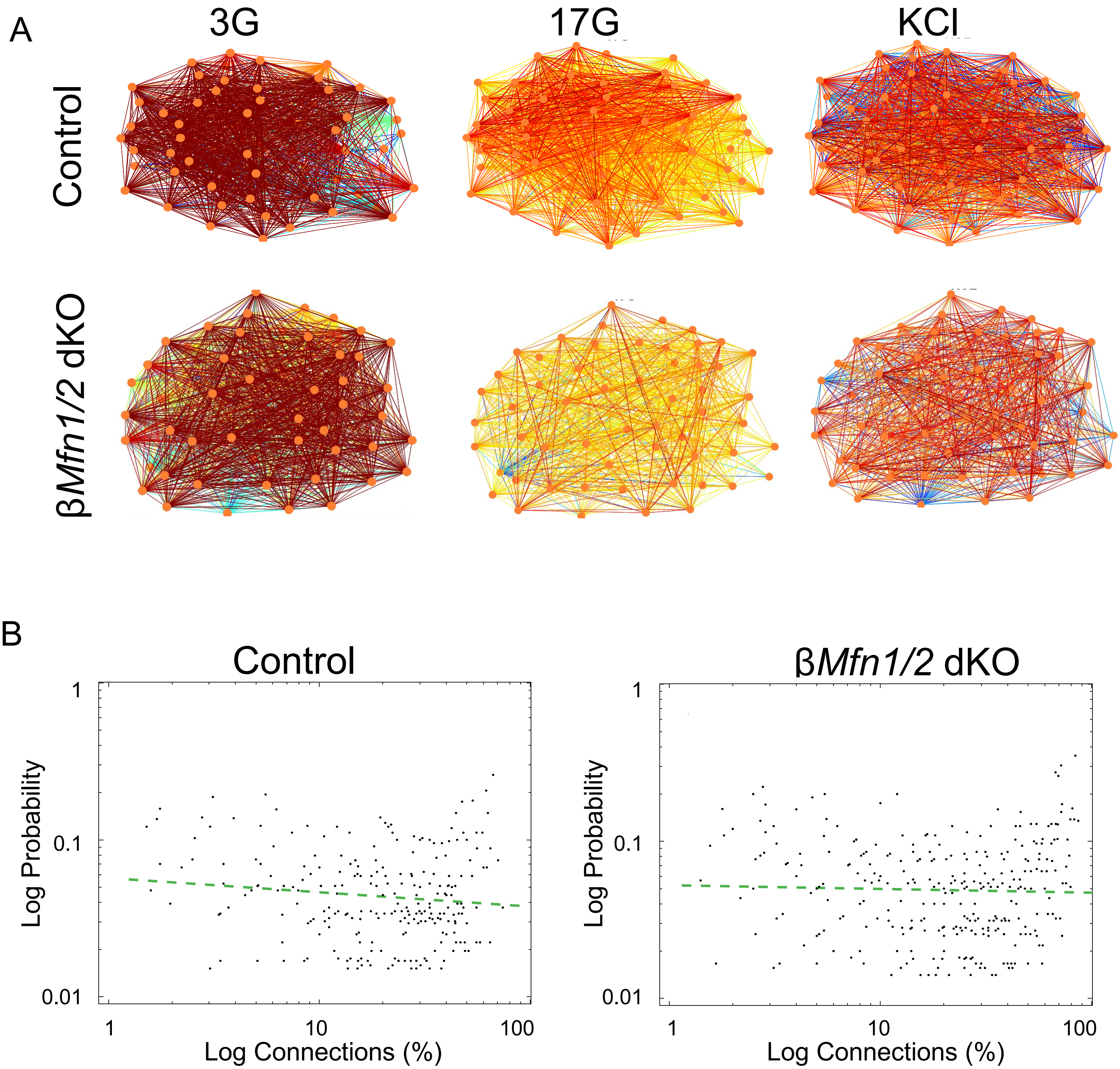
A
3G
17G
KCl
Control
βMfn1/2 dKO
B
Control
βMfn1/2 dKO
1
0.1
0.01
Log Probability
1
10
100
Log Connections (%)

Suppl. Fig.6

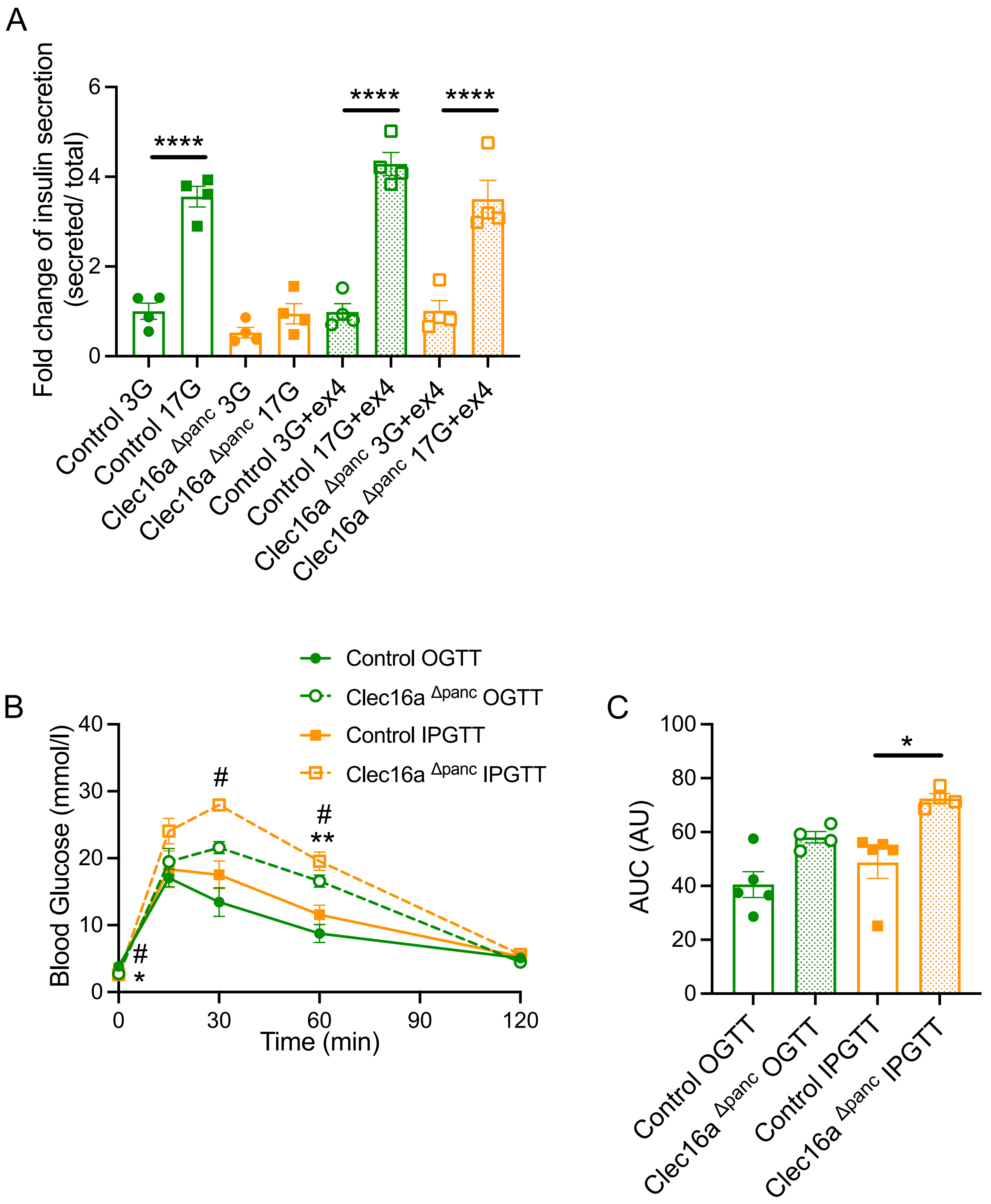
A
Fold change of insulin secretion (secreted/ total)
6
4
2
0
****
****
****
Control 3G
Control 17G
Clec16a Δpanc 3G
Clec16a Δpanc 17G
Control 3G+ex4
Control 17G+ex4
Clec16a Δpanc 3G+ex4
Clec16a Δpanc 17G+ex4
Control OGTT
Clec16a Δpanc OGTT
Control IPGTT
Clec16a Δpanc IPGTT
B
Blood Glucose (mmol/l)
40
30
20
10
0
#
#
**
#
*
0
30
60
90
120
Time (min)
C
AUC (AU)
100
80
60
40
20
0
*
Control OGTT
Clec16a Δpanc OGTT
Control IPGTT
Clec16a Δpanc IPGTT

Suppl. Fig.7

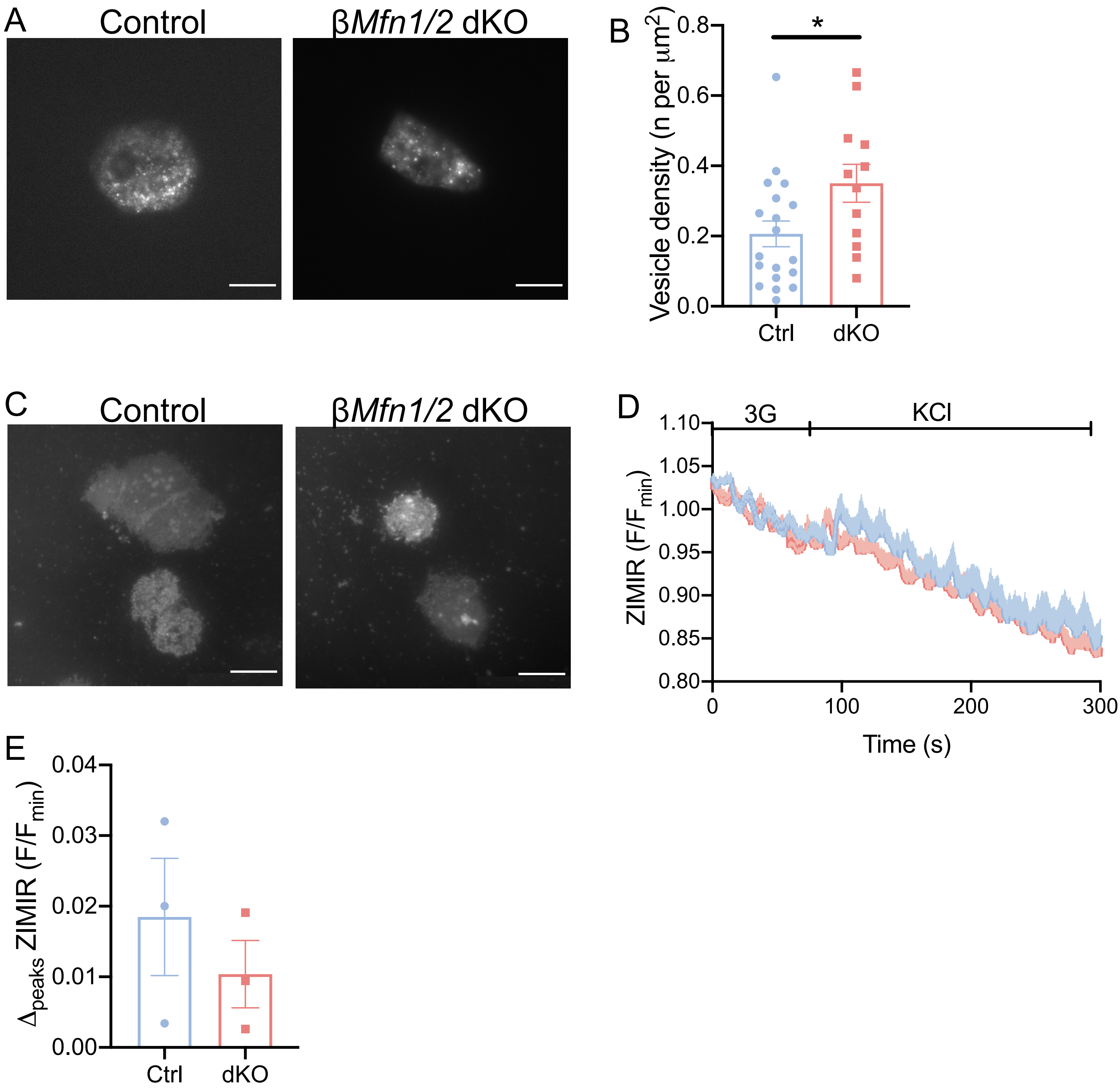
A
Control
βMfn1/2 dKO
B
*
Vesicle density (n per µm²)
0.0
0.2
0.4
0.6
0.8
Ctrl
dKO
C
Control
βMfn1/2 dKO
D
3G
KCl
ZIMIR (F/Fmin)
1.10
1.05
1.00
0.95
0.90
0.85
0.80
0
100
200
300
Time (s)
E
Δpeaks ZIMIR (F/Fmin)
0.04
0.03
0.02
0.01
0.00
Ctrl
dKO

Suppl. Fig.8

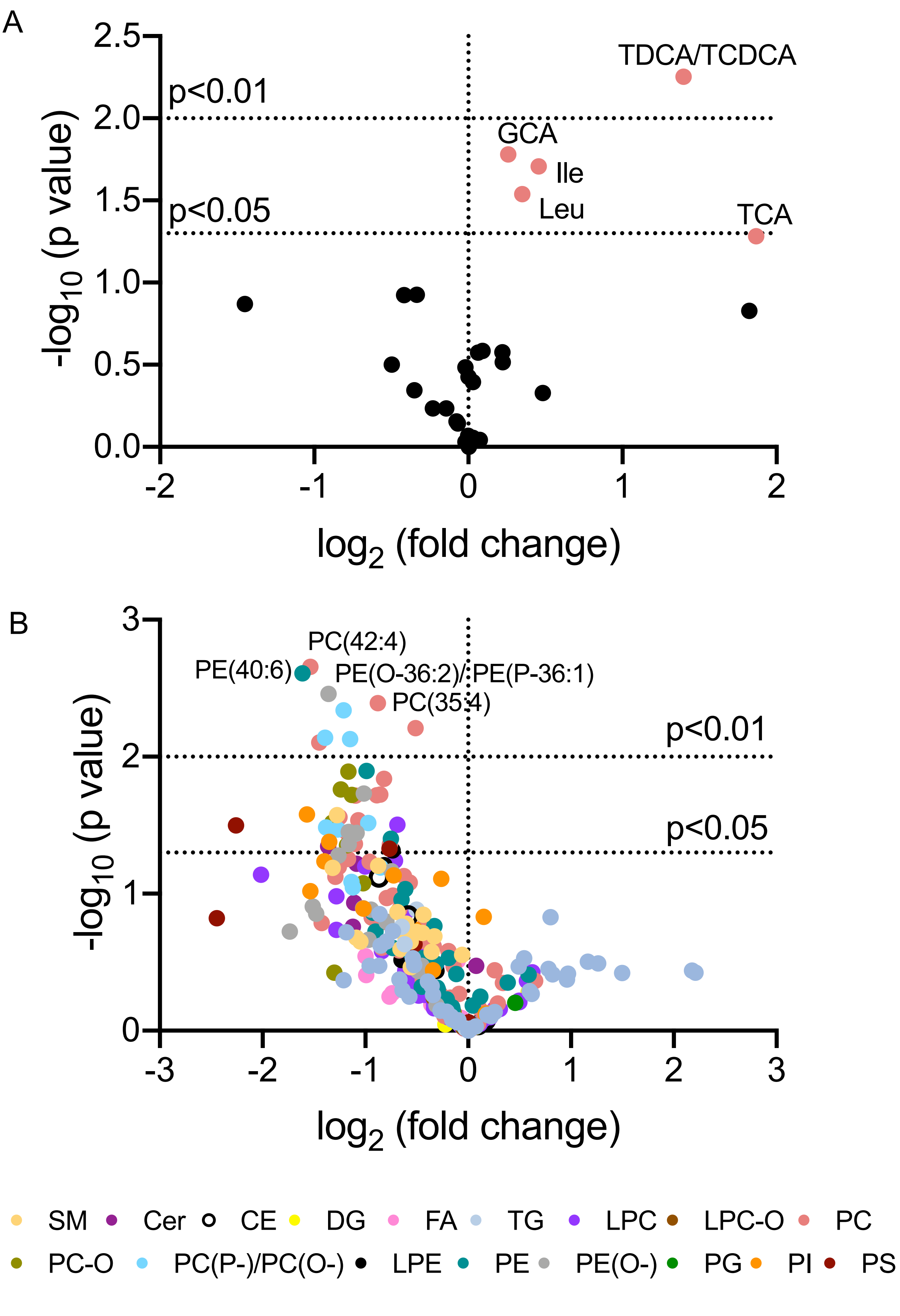
A
TDCA/TCDCA
p<0.01
GCA
Ile
Leu
p<0.05
TCA
-log10 (p value)
log2 (fold change)
B
PC(42:4)
PE(40:6)
PE(O-36:2)/PE(P-36:1)
PC(35:4)
p<0.01
p<0.05
SM
Cer
CE
DG
FA
TG
LPC
LPC-O
PC
PC-O
PC(P-)/PC(O-)
LPE
PE
PE(O-)
PG
PI
PS

Suppl. Fig.9

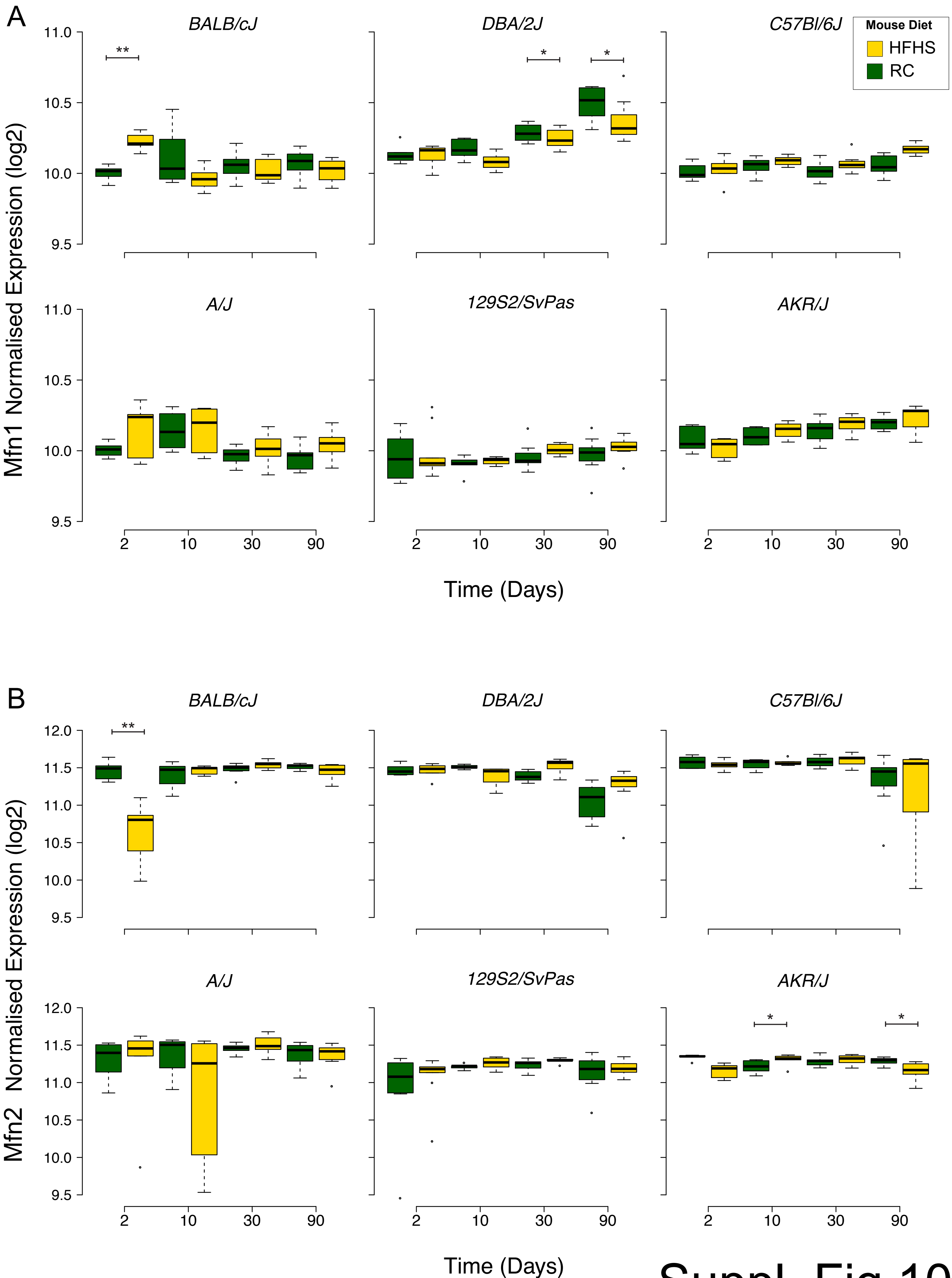
A
BALB/cJ
DBA/2J
C57Bl/6J
Mouse Diet
HFHS
RC
A/J
129S2/SvPas
AKR/J
Mfn1 Normalised Expression (log2)
Time (Days)
B
Mfn2 Normalised Expression (log2)
11.0
10.5
10.0
9.5
12.0
11.5
2
10
30
90
**
*


Suppl. Fig.10
